# Exact global alignment using A* with chaining seed heuristic and match pruning

**DOI:** 10.1101/2022.09.19.508631

**Authors:** Ragnar Groot Koerkamp, Pesho Ivanov

**Affiliations:** Department of Computer Science, ETH Zurich, Switzerland

## Abstract

**Motivation:** Sequence alignment has been at the core of computational biology for half a century. Still, it is an open problem to design a practical algorithm for exact alignment of a pair of related sequences in linear-like time (Medvedev, 2022b).

**Methods:** We solve exact global pairwise alignment with respect to edit distance by using the A* shortest path algorithm. In order to efficiently align long sequences with high divergence, we extend the recently proposed *seed heuristic* (Ivanov *et al*., 2022) with *match chaining, gap costs*, and *inexact matches*. We additionally integrate the novel *match pruning* technique and diagonal transition (Ukkonen, 1985) to improve the A* search. We prove the correctness of our algorithm, implement it in the A*PA aligner, and justify our extensions intuitively and empirically.

**Results:** On random sequences of divergence *d*=4% and length *n*, the empirical runtime of A*PA scales near-linearly with length (best fit *n*^1.06^, *n*≤10^7^ bp). A similar scaling remains up to *d*=12% (best fit *n*^1.24^, *n*≤10^7^ bp). For *n*=10^7^ bp and *d*=4%, A*PA reaches >500× speedup compared to the leading exact aligners EDLIB and BIWFA. The performance of A*PA is highly influenced by long gaps. On long (*n*>500 kbp) ONT reads of a human sample it efficiently aligns sequences with *d*<10%, leading to 3× median speedup compared to EDLIB and BIWFA. When the sequences come from different human samples, A*PA performs 1.7× faster than EDLIB and BIWFA.

**Availability:** github.com/RagnarGrootKoerkamp/astar-pairwise-aligner

## 1 Introduction

The problem of aligning one biological sequence to another is known as *global pairwise alignment* (Navarro, 2001). Among others, it is applied to genome assembly, read mapping, variant detection, and multiple sequence alignment (Prjibelski *et al*., 2019). Despite the centrality and age of pairwise alignment (Needleman and Wunsch, 1970), “a major open problem is to implement an algorithm with linear-like empirical scaling on inputs where the edit distance is linear in *n*” (Medvedev, 2022b).

Alignment accuracy affects subsequent analyses, so a common goal is to find a shortest sequence of edit operations (single letter insertions, deletions, and substitutions) that transforms one sequence into the other. The length of such a sequence is known as *Levenshtein distance* (Levenshtein, 1966) and *edit distance*. It has recently been proven that edit distance can not be computed in strongly subquadratic time, unless SETH is false (Backurs and Indyk, 2015). When the number of sequencing errors is proportional to the length, existing exact aligners scale quadratically both in the theoretical worst case and in practice. Given the increasing amounts of biological data and increasing read lengths, this is a computational bottleneck (Kucherov, 2019).

We solve the global alignment problem provably correct and empirically fast by using A* on the alignment graph and building on many existing techniques. Our implementation A*PA (A* Pairwise Aligner) scales near-linear with length up to 10^7^ bp long sequences with divergence up to 12%. Additionally, it shows a speedup over other highly optimized aligners when aligning long ONT reads.

### 1.1 Overview of method

To align two sequences *A* and *B* globally with minimal cost, we use the A* shortest path algorithm from the start to the end of the alignment graph, as first suggested by Hadlock (1988b). A core part of the A* algorithm is the heuristic function *h*(*u*) that provides a lower bound on the remaining distance from the current vertex *u*. A good heuristic efficiently computes an accurate estimate *h*, so suboptimal paths get penalized more and A* prioritizes vertices on a shortest path, thus reaching the target quicker. In this paper, we extend the *seed heuristic* by Ivanov *et al*. (2022) in several ways to increase its accuracy for long and erroneous sequences.

#### Seed heuristic (SH)

To define the *seed heuristic h*s, we split *A* into short, non-overlapping substrings (*seeds*) of fixed length *k* (Fig. 2a). Since the whole sequence *A* has to be aligned, each of the seeds also has to be aligned somewhere in *B*. If a seed does not match anywhere in *B* without mistakes, then at least 1 edit has to be made to align it. Thus, the seed heuristic *h*_s_ is the number of remaining seeds (contained in *A*_≥*i*_) that do not match anywhere in *B*. The seed heuristic is a lower bound on the distance between the remaining suffixes *A*_≥*i*_ and *B*_≥*j*_. In order to compute *h*s efficiently, we precompute all *matches* in *B* for all seeds from *A*. Where Ivanov *et al*. (2022) uses *crumbs* to mark upcoming matches in the graph, we do not need them due to the simpler structure of sequence-to-sequence alignment.

**Fig. 1.**
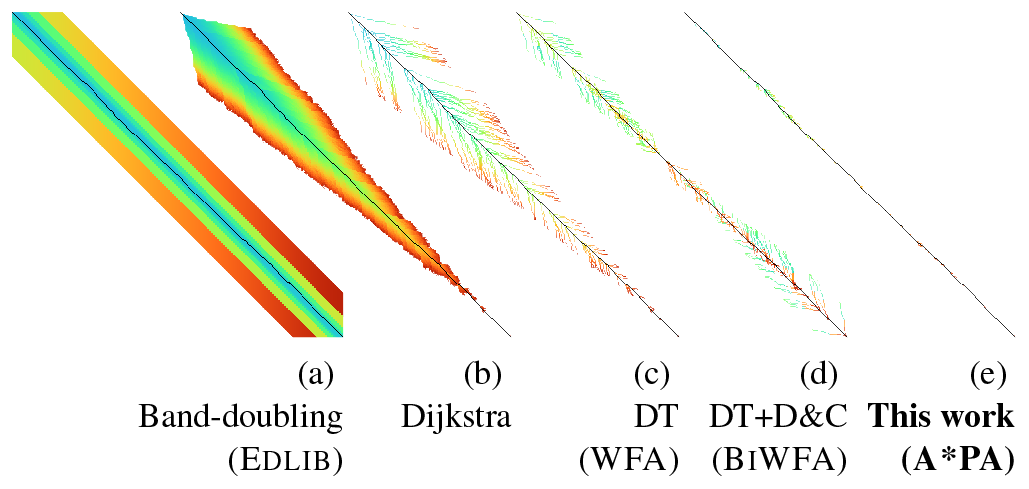
Computed states per algorithm. Various optimal alignment algorithms and their implementation are demonstrated on synthetic data (length *n*=500 bp, divergence *d*=16%). The colour indicates the order of computation from blue to red. (a) Band-doubling (E_DLIB_), (b) Dijkstra, (c) Diagonal transition/DT (WFA), (d) DT with divide-and-conquer/D&C (B_I_WFA), (e) A*PA with gap-chaining seed heuristic (GCSH), match pruning, and DT (seed length *k*=5 and exact matches).

**Fig. 2.**
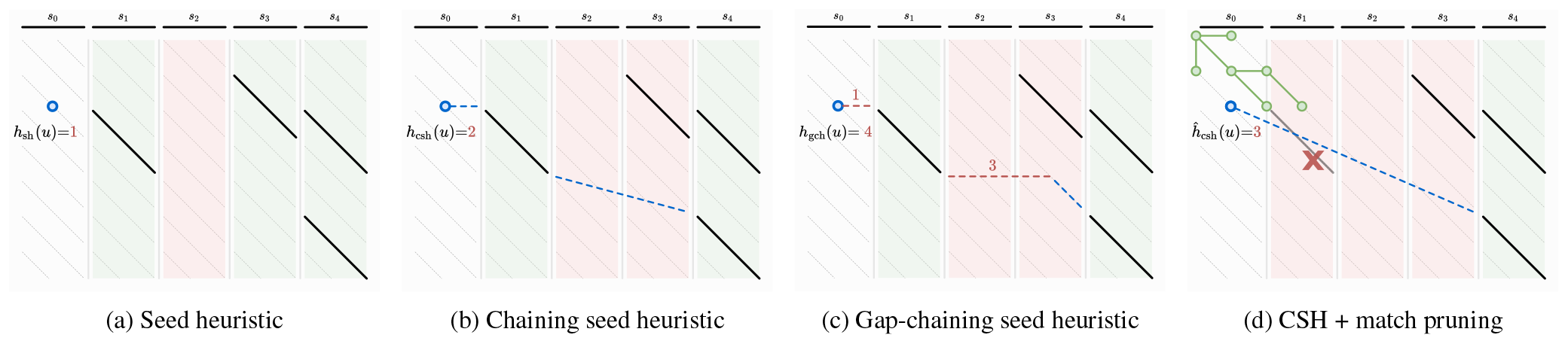
Demonstration of seed heuristic, chaining seed heuristic, gap-chaining seed heuristic, and match pruning. Sequence *A* on top is split into 5 seeds (horizontal black segments). Each seed is exactly matched in *B* (diagonal black segments). The heuristic is evaluated at state *u* (blue circles 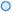), based on the 4 remaining seeds. The heuristic value is based on a maximal chain of matches (green columns 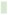 for seeds with matches; red columns 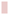 otherwise). Dashed lines denote chaining of matches. (a) The seed heuristic *h*_s_(*u*)=1 is the number of remaining seeds that do not have matches (only *s*_2_). (b) The chaining seed heuristic *h*_cs_(*u*)=2 is the number of remaining seeds without a match (*s*_2_ and *s*_3_) on a path going only down and to the right containing a maximal number of matches. (c) The gap-chaining seed heuristic *h*_gcs_(*u*)=4 is minimal cost of a chain, where the cost of joining two matches is the maximum of the number of not matched seeds and the gap cost between them. Red dashed lines denote gap costs. (d) Once the start or end of a match is expanded (green circles 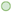), the match is *pruned* (red cross 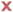), and future computations of the heuristic ignore it. *s*_1_ is removed from the maximum chain of matches starting at *u* so *ĥ*_*cs*_(*u*) increases by 1.

#### Chaining (CSH)

One drawback of the SH is that it may use matches that do not lie together on a path from *u* to the end, as for example the matches for *s*_1_ and *s*_3_ in Fig. 2a. In the *chaining seed heuristic h*cs (Sec. 3.1), we enforce that the matches occur in the same order in *B* as their corresponding seeds occur in *A*, i.e., the matches form a *chain* going down and right (Fig. 2b). Now, the number of upcoming errors is at least the minimal number of remaining seeds that cannot be aligned on a single chain to the target. When there are many spurious matches (i.e. outside the optimal alignment), chaining improves the accuracy of the heuristic, thus reducing the number of states expanded by A*. To compute CSH efficiently, we subtract the maximal number of matches in a chain starting in the current state from the number of remaining seeds.

#### Gap costs (GCSH)

The CSH penalizes the chaining of two matches by the *seed cost*, the number of skipped seeds in between them. This chaining may skip a different number of letters in *A* and *B*, in which case the absolute difference between these lengths (*gap cost*) is a lower bound on the length of a path between the two matches. The *gap-chaining seed heuristic h*gcs (Fig. 2c) takes the maximum of the gap cost and the seed cost, which significantly improves the accuracy of the heuristic for sequences with long indels.

#### Inexact matches

To further improve the accuracy of the heuristic for divergent sequences, we use *inexact matches* (Wu and Manber, 1992; Marco-Sola *et al*., 2012). For each seed in *A*, our algorithm now finds all its inexact matches in *B* with cost at most 1. The lack of a match of a seed then implies that at least *r*=2 edits are needed to align it. This doubles the *potential* of our heuristic to penalize errors.

#### Match pruning

In order to further improve the accuracy of our heuristic, we apply the *multiple-path pruning* observation (Poole and Mackworth, 2017): once a shortest path to a vertex *u* has been found, no other path to *u* can be shorter. Since we search for a single shortest path, we want to incrementally update our heuristic (similar to Real-Time Adaptive A* (Koenig and Likhachev, 2006)) to penalize further paths to *u*. We prove that once A* expands a state *u* which is at the start or end of a match, indeed it has found a shortest path to *u*. Then we can ignore (*prune*) such a match, thus penalizing other paths to *u* (Fig. 2d, Sec. 3.2). Pruning increases the heuristic in states preceding the match, thereby penalizing states preceding the “tip” of the A* search. This reduces the number of expanded states, and leads to near-linear scaling with sequence length (Fig. 1e).

#### Diagonal transition (DT)

The diagonal transition algorithm only visits so called *farthest reaching* states (Ukkonen, 1985; Myers, 1986) along each diagonal and lies at the core of WFA (Marco-Sola *et al*., 2021) (Fig. 1c). We introduce the *diagonal transition* optimization to the A* algorithm that skips states known to be not farthest reaching. This is independent of the A* heuristic and makes the exploration more “hollow”, especially speeding up the quadratic behavior of A* in complex regions.

We present an algorithm to efficiently initialize and evaluate these heuristics and optimizations (Sec. 3.3 and App. A), prove the correctness of our methods (App. B), and evaluate and compare their performance to other optimal aligners (Sec. 4 and App. C).

### 1.2 Related work

We first outline the algorithms behind the fastest exact global aligners: DP-based band doubling (used by EDLIB) and diagonal transition (used by BIWFA). Then, we outline methods that A*PA integrates.

#### Dynamic programming (DP)

This classic approach to aligning two sequences computes a table where each cell contains the edit distance between a prefix of the first sequence and a prefix of the second by reusing the solutions for shorter prefixes. This quadratic DP was introduced for speech signals Vintsyuk (1968) and genetic sequences (Needleman and Wunsch, 1970; Sankoff, 1972; Sellers, 1974; Wagner and Fischer, 1974). The quadratic *O*(*nm*) runtime for sequences of lengths *n* and *m* allowed for aligning of long sequences for the time but speeding it up has been a central goal in later works. Implementations of this algorithm include SEQAN (Reinert *et al*., 2017) and PARASAIL (Daily, 2016).

#### Band doubling and bit-parallelization

When the aligned sequences are similar, the whole DP table does not need to be computed. One such output-sensitive algorithm is the *band doubling* algorithm of Ukkonen (1985) (Fig. 1a) which considers only states around the main diagonal of the table, in a *band* with exponentially increasing width, leading to *O*(*ns*) runtime, where *s* is the edit distance between the sequences. This algorithm, combined with the *bit-parallel optimization* by Myers (1999) is implemented in EDLIB (Šošić and Šikić, 2017) with *O*(*ns*/*w*) runtime, where *w* is the machine word size (nowadays 64).

#### Diagonal transition (DT)

The *diagonal transition* algorithm (Ukkonen, 1985; Myers, 1986) exploits the observation that the edit distance does not decrease along diagonals of the DP matrix. This allows for an equivalent representation of the DP table based on *farthest-reaching states* for a given edit distance along each diagonal. Diagonal transition has an *O*(*ns*) worst-case runtime but only takes expected *O*(*n*+*s*^2^) time (Fig. 1c) for random input sequences (Myers, 1986) (which is still quadratic for a fixed divergence *d* = *s*/*n*). It has been extended to linear and affine costs in the *wavefront alignment* (WFA) (Marco-Sola *et al*., 2021) in a way similar to Gotoh (1982). Its memory usage has been improved to linear in BIWFA (Marco-Sola *et al*., 2023) by combining it with the divide-and-conquer approach of Hirschberg (1975), similar to Myers (1986) for unit edit costs. Wu *et al*. (1990) and Papamichail and Papamichail (2009) apply diagonal transition to align sequences of different lengths.

#### Contours

The longest common subsequence (LCS) problem is a special case of edit distance, in which gaps are allowed but substitutions are forbidden. *Contours* partition the state-space into regions with the same remaining answer of the LCS subtask (Fig. 3). The contours can be computed in log-linear time in the number of matching elements between the two sequences which is practical for large alphabets (Hirschberg, 1977; Hunt and Szymanski, 1977).

**Fig. 3.**
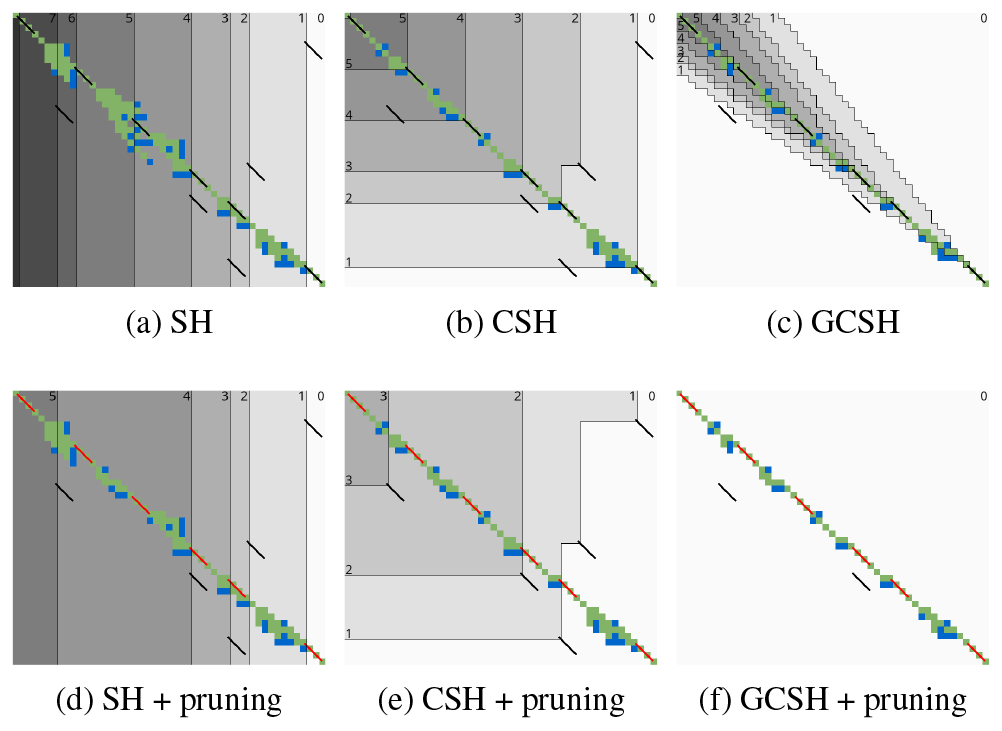
Contours and layers of different heuristics after aligning. (*n*=48, *m*=42, *r*=1, *k*=3, edit distance 10). Exact matches are black diagonal segments 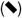. The background colour indicates *Sp*(*u*), the maximum number of matches on a ⪯_*p*_-chain from *u* to the end starting, with *Sp*(*u*) = 0 in white. The thin black boundaries of these regions are *Contours*. The states of layer ℒ_*ℓ*_ *precede* contour *ℓ*. Expanded states are green 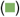, open states blue 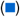, and pruned matches red 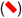. Pruning matches changes the contours and layers. GCSH ignores matches 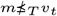.

#### Shortest paths and A*

An alignment that minimizes edit distance corresponds to a shortest path in the *alignment graph* (Vintsyuk, 1968; Ukkonen, 1985). Assuming non-negative edit costs, a shortest path can be found using Dijkstra’s algorithm (Ukkonen, 1985) (Fig. 1b) or A* (Hart *et al*., 1968). A* is an informed search algorithm which uses a task-specific heuristic function to direct its search, and has previously been applied to the alignment graph by Hadlock (1988a,b) and Spouge (1989, 1991). A* with an accurate heuristic may find a shortest path significantly faster than an uninformed search such as Dijkstra’s algorithm.

#### A* heuristics

One widely used heuristic function is the *gap cost* that counts the minimal number of indels needed to align the suffixes of two sequences (Ukkonen, 1985; Myers and Miller, 1995; Spouge, 1989; Wu *et al*., 1990; Papamichail and Papamichail, 2009; Šošić and Šikić, 2017). Hadlock (1988b) introduces a heuristic based on character frequencies.

#### Seed-and-extend

*Seed-and-extend* is a commonly used paradigm for approximately solving semi-global alignment by first matching similar regions between sequences (*seeding*) to find *matches* (also called *anchors*), followed by *extending* these matches (Kucherov, 2019). Aligning long reads requires the additional step of chaining the seed matches (*seed-chain-extend*). Seeds have also been used to solve the LCSk generalization of LCS (Benson *et al*., 2014; Pavetić *et al*., 2017). Except for the seed heuristic (Ivanov *et al*., 2022), most seeding approaches seek for seeds with accurate long matches.

#### Seed heuristic

A* with *seed heuristic* is an exact algorithm that was recently introduced for exact semi-global sequence-to-graph alignment (Ivanov *et al*., 2022). In a precomputation step, the query sequence is split into non-overlapping *seeds* each of which is matched exactly to the reference. When A* explores a new state, the seed heuristic is computed as the number of remaining seeds that cannot be matched in the upcoming reference. A* with the seed heuristic enables provably-exact alignment but runs reasonably-fast only when the long sequences are very similar (≤ 0.3% divergence).

### 1.3 Contributions

We present an algorithm for exact global alignment that uses A* on the alignment graph (Hart *et al*., 1968; Hadlock, 1988b), starting with the seed heuristic of Ivanov *et al*. (2022).

We increase the accuracy of this heuristic in several novel ways: seeds must match in order in the *chaining seed heuristic*, and gaps between seeds are penalized in the *gap-chaining seed heuristic*. The novel *match pruning* technique penalizes states “lagging behind” the tip of the search and turns the otherwise quadratic algorithm into an empirically near-linear algorithm in many cases. Inexact matches (Wu and Manber, 1992; Marco-Sola *et al*., 2012) increase the divergence of sequences that can be efficiently aligned. We additionally apply the diagonal transition algorithm (Ukkonen, 1985; Myers, 1986), so that only the small fraction of farthest-reaching states needs to be computed. We prove the correctness of our methods, and apply contours (Hirschberg, 1977; Hunt and Szymanski, 1977) to efficiently initialize and evaluate the heuristic. We implement our method in the novel aligner A*PA.

On uniform random synthetic data with 4% divergence, the runtime of A*PA scales linearly with length up to 10^7^ bp and is up to 500× faster than EDLIB and BIWFA. On >500 kbp long Oxford Nanopore (ONT) reads of the human genome, A*PA is 3× faster in median than EDLIB and BIWFA when only read errors are present, and 1.7× faster in median when additionally genetic variation is present.

## 2 Preliminaries

This section provides definitions and notation that are used throughout the paper. A summary of notation is included in App. D.

### Sequences

The input sequences 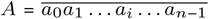 and 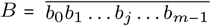 are over an alphabet Σ with 4 letters. We refer to substrings 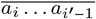 as *A*_*i*…*i*′_, to prefixes 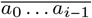 as *A*_<*i*_, and to suffixes 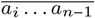 as *A*_≥*i*_. The *edit distance* ed(*A, B*) is the minimum number of insertions, deletions, and substitutions of single letters needed to convert *A* into *B*. The *divergence* is the observed number of errors per letter, *d ≔*ed(*A, B*)/*n*, whereas the *error rate e* is the number of errors per letter *applied* to a sequence.

### Alignment graph

Let *state* ⟨*i, j*⟩ denote the subtask of aligning the prefix *A*_<*i*_ to the prefix *B*_<*j*_. The *alignment graph* (also called *edit graph*) *G*(*V, E*) is a weighted directed graph with vertices *V* = {⟨*i, j*⟩ ∣ 0 ≤ *i* ≤ *n*, 0 ≤ *j* ≤ *m*} corresponding to all states, and edges connecting subtasks: edge ⟨*i, j*⟩ → ⟨*i*+1, *j*+1⟩ has cost 0 if *a*_*i*_ = *b*_*j*_ (match) and 1 otherwise (substitution), and edges ⟨*i, j*⟩ → ⟨*i*+1, *j*⟩ (deletion) and ⟨*i, j*⟩ → ⟨*i, j*+1⟩ (insertion) have cost 1. We denote the starting state ⟨0, 0⟩ by *vs*, the target state ⟨*n, m*⟩ by *vt*, and the distance between states *u* and *v* by d(*u, v*). For brevity we write *f* ⟨*i, j*⟩ instead of *f* (⟨*i, j*⟩).

### Paths and alignments

A path *π* from ⟨*i, j*⟩ to ⟨*i*^′^, *j*^′^⟩ in the alignment graph *G* corresponds to a *(pairwise) alignment* of the substrings *A*_*i*…*i*′_ and *B*_*j*…*j*′_ with cost c_path_(*π*). A shortest path *π*^∗^ from *v*_*s*_ to *v*_*t*_ corresponds to an optimal alignment, thus c_path_(*π*^∗^) = d(*vs, vt*) = ed(*A, B*). We write *g*^∗^(*u*) ≔ d(*vs, u*) for the distance from the start to *u* and *h*^∗^(*u*) ≔d(*u, vt*) for the distance from *u* to the target.

### Seeds and matches

We split the sequence *A* into a set of consecutive non-overlapping substrings (*seeds*) 𝒮 = {*s*_0_, *s*_1_, *s*_2_, …, *s*_⌊*n*/*k*⌋−1_}, such that each seed *s*_*l*_=*A*_*lk*…*lk*+*k*_ has length *k*. After aligning the first *i* letters of *A*, our heuristics will only depend on the *remaining* seeds 𝒮_≥*i*_ *≔* {*s*_*l*_ ∈ 𝒮 ∣ *lk* ≥ *i*} contained in the suffix *A*_≥*i*_. We denote the set of seeds between *u*=⟨*i, j*⟩ and *v*=⟨*i*^′^, *j*^′^⟩ by 𝒮_*u*…*v*_ = 𝒮_*i*…*i*_′ = {*s*_*l*_ ∈ 𝒮 ∣ *i* ≤ *lk, lk* + *k* ≤ *i*^′^} and an *alignment* of *s* to a subsequence of *B* by *π*_*s*_. The alignments of seed *s* with sufficiently low cost (Sec. 3.1) form the set ℳ_*s*_ of *matches*.

### Dijkstra and A*

Dijkstra’s algorithm (Dijkstra, 1959) finds a shortest path from *vs* to *vt* by *expanding* (generating all successors) vertices in order of increasing distance *g*^∗^(*u*) from the start. Each vertex to be expanded is chosen from a set of *open* vertices. The A* algorithm (Hart *et al*., 1968, 1972; Pearl, 1984), instead directs the search towards a target by expanding vertices in order of increasing *f* (*u*) ≔ *g*(*u*) + *h*(*u*), where *h*(*u*) is a heuristic function that estimates the distance *h*^∗^(*u*) to the end and *g*(*u*) is the shortest length of a path from *vs* to *u* found so far. A heuristic is *admissible* if it is a lower bound on the remaining distance, *h*(*u*) ≤ *h*^∗^(*u*), which guarantees that A* has found a shortest path as soon as it expands *v*_*t*_. Heuristic *h*_1_ *dominates* (is *more accurate* than) another heuristic *h*_2_ when *h*_1_(*u*) ≥ *h*_2_(*u*) for all vertices *u*. A dominant heuristic will usually, but not always (Holte, 2010), expand less vertices. Note that Dijkstra’s algorithm is equivalent to A* using a heuristic that is always 0, and that both algorithms require non-negative edge costs. Our variant of the A* algorithm is provided in App. A.1.

### Chains

A state *u* = ⟨*i, j*⟩ ∈ *V precedes* a state *v* = ⟨*i*^′^, *j*^′^⟩ ∈ *V*, denoted *u* ⪯ *v*, when *i* ≤ *i*^′^ and *j* ≤ *j*^′^. Similarly, a match *m* precedes a match *m*^′^, denoted *m* ⪯ *m*^′^, when the end of *m* precedes the start of *m*^′^. This makes the set of matches a partially ordered set. A state *u* precedes a match *m*, denoted *u* ⪯ *m*, when it precedes the start of the match. A *chain* of matches is a (possibly empty) sequence of matches *m*_1_ ⪯ ⃛ ⪯ *m*_*l*_.

### Gap cost

The number of indels to align substrings *A*_*i*…*i*′_ and *B*_*j*…*j*′_ is at least their difference in length: *c*_gap_(⟨*i, j*⟩, ⟨*i*^′^, *j*^′^⟩) ≔ ∣(*i*^′^−*i*)−(*j*^′^−*j*)∣. For *u* ⪯ *v* ⪯ *w*, the gap cost satisfies the triangle inequality *c*_gap_(*u, w*) ≤ *c*_gap_(*u, v*) + *c*_gap_(*v, w*).

### Contours

To efficiently calculate maximal chains of matches, *contours* are used. Given a set of matches ℳ, *S*(*u*) is the number of matches in the longest chain *u* ⪯ *m*_1_ ⪯ …, starting at *u*. The function *S*⟨*i, j*⟩ is non-increasing in both *i* and *j. Contours* are the boundaries between regions of states with *S*(*u*) = *ℓ* and *S*(*u*) < *ℓ* (Fig. 3). Note that contour *ℓ* is completely determined by the set of matches *m* ∈ ℳ for which *S*(start(*m*)) = *ℓ* (Hirschberg, 1977). Hunt and Szymanski (1977) give an algorithm to efficiently compute *S* when ℳ is the set of single-letter matches between *A* and *B*, and Deorowicz and Grabowski (2014) give an algorithm when ℳ is the set of exact *k*-mer matches.

## 3 Methods

We formally define the general chaining seed heuristic (Sec. 3.1) that encompases *inexact matches, chaining*, and *gap costs* (Fig. 2). Next, we introduce the *match pruning* (Sec. 3.2) improvement and integrate our A* algorithm with the *diagonal-transition* optimization (App. A.2). We present a practical algorithm (Sec. 3.3), implementation (App. A.3) and proofs of correctness (App. B).

### 3.1 General chaining seed heuristic

We introduce three heuristics for A* that estimate the edit distance between a pair of suffixes. Each heuristic is an instance of a *general chaining seed heuristic*. After splitting the first sequence into seeds S, and finding all matches ℳ in the second sequence, any shortest path to the target can be partitioned into a *chain* of matches and connections between the matches. Thus, the cost of a path is the sum of match costs cm and *chaining costs γ*. Our simplest seed heuristic ignores the position in *B* where seeds match and counts the number of seeds that were not matched (*γ*=*c*_seed_). To efficiently handle more errors, we allow seeds to be matched inexactly, require the matches in a path to be ordered (CSH), and include the gap-cost in the chaining cost *γ*= max(*c*_gap_, *c*_seed_) to penalize indels between matches (GCSH).

#### Inexact matches

We generalize the notion of exact matches to *inexact matches*. We fix a threshold cost *r* (0<*r*≤*k*) called the *seed potential* and define the set of *matches ℳ*_*s*_ as all alignments *m* of seed *s* with *match cost* cm(*m*) < *r*. The inequality is strict so that ℳ_*s*_ = ∅ implies that aligning the seed will incur cost at least *r*. Let ℳ = ⋃_*s*_ *ℳ*_*s*_ denote the set of all matches. With *r*=1 we allow only *exact* matches, while with *r*=2 we allow both exact and *inexact* matches with one edit. We do not consider higher *r* in this paper. For notational convenience, we define *mω* ∉ ℳ to be a match from *vt* to *vt* of cost 0.

#### Potential of a heuristic

We call the maximal value the heuristic can take in a state its *potential P*. The potential of our heuristics in state ⟨*i, j*⟩ is the sum of seed potentials *r* over all seeds after *i*: *P* ⟨*i, j*⟩ ≔ *r* ⋅∣𝒮_≥*i*_∣.

#### Chaining matches

Each heuristic depends on a *partial order* on states that limits how matches can be *chained*. We write *u* ⪯*p v* for the partial order implied by a function *p*: *p*(*u*) ⪯ *p*(*v*). A ⪯*p-chain* is a sequence of matches *m*_1_ ⪯_*p*_ ⋅⋅⋅⪯_*p*_ *m*_*l*_ that precede each other: end(*m*_*i*_) ⪯_*p*_ start(*m*_*i*+1_) for 1 ≤ *i* < *l*. To chain matches according only to their *i*-coordinate, SH is defined using ⪯_*i*_-chains, while CSH and GCSH are defined using ⪯ that compares both *i* and *j*.

#### Chaining cost

The *chaining cost γ* is a lower bound on the path cost between two consecutive matches: from the end state *u* of a match, to the start *v* of the next match.

For SH and CSH, the *seed cost* is *r* for each seed that is not matched: *c*_seed_(*u, v*) ≔ *r* ⋅∣𝒮_*u*…*v*_∣. When *u* ⪯_*i*_ *v* and *v* is not in the interior of a seed, then *c*_seed_(*u, v*) = *P* (*u*) − *P* (*v*).

For GCSH, we also include the gap cost *c*gap(⟨*i, j*⟩, ⟨*i*^′^, *j*^′^⟩) ≔ ∣(*i*^′^−*i*)−(*j*^′^−*j*)∣ which is the minimal number of indels needed to correct for the difference in length between the substrings *Ai*…*i*′ and *Bj*…*j*′ between two consecutive matches (Sec. 2). Combining the seed cost and the chaining cost, we obtain the gap-seed cost *c*_gs_ = max(*c*_seed_, *c*_gap_), which is capable of penalizing long indels and we use for GCSH. Note that *γ*=*c*_seed_+*c*_gap_ would not give an admissible heuristic since indels could be counted twice, in both *c*_seed_ and *c*_gap_.

For conciseness, we also define *γ, c*_seed_, *c*_gap_, and *c*_gs_ between matches *γ*(*m, m*^′^) ≔ *γ*(end(*m*), start(*m*^′^)), from a state to a match *γ*(*u, m*^′^) ≔ *γ*(*u*, start(*m*^′^)), and from a match to a state *γ*(*m, u*) = *γ*(end(*m*), *u*).

#### General chaining seed heuristic

We now define the general chaining seed heuristic that we use to instantiate SH, CSH and GCSH.

##### Definition 1

(General chaining seed heuristic). *Given a set of matches ℳ, partial order* ⪯_*p*_, *and chaining cost γ, the* general chaining seed heuristic 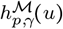 *is the minimal sum of match costs and chaining costs over all* ⪯_*p*_*-chains (indexing extends to m*_0_ ≔ *u and m*_*l*+1_ ≔ *m*_*ω*_):

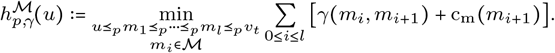

We instantiate our heuristics according to Table 1. Our admissibility proofs (App. B.1) are based on cm and *γ* being lower bounds on disjoint parts of the remaining path. Since the more complex *h*_gcs_ dominates the other heuristics it usually expand fewer states.

**Table 1.** Definitions of our heuristic functions. SH orders the matches by *i* and uses only the seed cost. CSH orders the matches by both *i* and *j*. GCSH additionally exploits the gap cost.

| Heuristic | Order | Chaining cost $\gamma$ |
| --- | --- | --- |
| $h_s(u)$ Seed heuristic (SH) | $\leq_i$ | $c_{\text{seed}}$ |
| $h_{\text{cs}}(u)$ Chaining seed h. (CSH) | ≤ | $c_{\text{seed}}$ |
| $h_{\text{gcs}}(u)$ Gap-chaining seed h. (GCSH) | ≤ | $\max(c_{\text{gap}}, c_{\text{seed}})$ |

##### Theorem 1

*The seed heuristic h_s_, the chaining seed heuristic h_cs_, and the gap-chaining seed heuristic h_gcs_ are admissible. Furthermore, 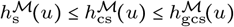 for all states u.*

We are now ready to instantiate A* with our admissible heuristics but we will first improve them and show how to compute them efficiently.

### 3.2 Match pruning

In order to reduce the number of states expanded by the A* algorithm, we apply the *multiple-path pruning* observation: once a shortest path to a state has been found, no other path to this state could possibly improve the global shortest path (Poole and Mackworth, 2017). As soon as A* expands the start or end of a match, we *prune* it, so that the heuristic in preceding states no longer benefits from the match, and they get deprioritized by A*. We define *pruned* variants of all our heuristics that ignore pruned matches:

#### Definition 2

(Pruning heuristic). *Let E be the set of expanded states during the A* search, and let ℳ*/*E be the set of matches that were not pruned, i*.*e. those matches not starting or ending in an expanded state. We say that ĥ* ≔ *h*^*ℳ*/*E*^ *is a* pruning heuristic *version of h*.

The hat over the heuristic function (ĥ) denotes the implicit dependency on the progress of the A*, where at each step a different *h*^*ℳ*/*E*^ is used. Our modified A* algorithm (App. A.1) works for pruning heuristics by ensuring that the *f* -value of a state is up-to-date before expanding it, and otherwise *reorders* it in the priority queue. Even though match pruning violates the admissibility of our heuristics for some vertices, we prove that A* is sill guaranteed to find a shortest path (App. B.2). To this end, we show that our pruning heuristics are *weakly-admissible heuristics* (Def. 7) in the sense that they are admissible on at least one path from *vs* to *vt*.

#### Theorem 2

*A* with a weakly-admissible heuristic finds a shortest path*.

#### Theorem 3

*The pruning heuristics ĥ_s_, ĥ_cs_, ĥ_gcs_ are weakly admissible.*

Pruning will allow us to scale near-linearly with sequence length, without sacrificing optimality of the resulting alignment.

### 3.3 Computing the heuristic

We present an algorithm to efficiently compute our heuristics (pseudocode in App. A.4, worst-case asymptotic analysis in App. A.5). At a high level, we rephrase the minimization of costs (over paths) to a maximization of *scores* (over chains of matches). We initialize the heuristic by precomputing all seeds, matches, potentials and a *contours* data structure used to compute the maximum number of matches on a chain. During the A* search, the heuristic is evaluated in all explored states, and the contours are updated whenever a match gets pruned.

#### Scores

The *score of a match m* is score(*m*)≔*r*− c_m_(*m*) and is always positive. The *score of a* ⪯_*p*_*-chain m*_1_ ⪯_*p*_ ⋅⋅⋅⪯_*p*_ *m*_*l*_ is the sum of the scores of the matches in the chain. We define the chain score of a match *m* as

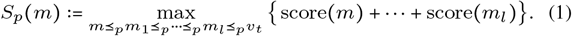

Since ⪯_*p*_ is a partial order, *S*_*p*_ can be computed with base case *S*_*p*_(*m*_*ω*_) = 0 and the recursion

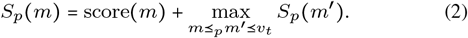

We also define the chain score of a state *u* as the maximum chain score over succeeding matches 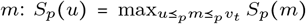, so that Eq. (2) can be rewritten as *S*_*p*_(*m*) = score(*m*) + *S*_*p*_(end(*m*)).

The following theorem allows us to rephrase the heuristic in terms of potentials and scores for heuristics that use *γ*=*c*_seed_ and respect the order of the seeds, which is the case for *h*s and *h*cs (proof in App. B.3):

##### Theorem 4

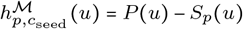 *for any partial order* ⪯_*p*_ *that is a refinement of* ⪯_*i*_ *(i*.*e. u* ⪯_*p*_ *v must imply u* ⪯_*i*_ *v)*.

#### Layers and contours

We compute *h*_s_ and *h*_cs_ efficiently using *contours*. Let *layer ℒ*_*ℓ*_ be the set of states *u* with score *S*_*p*_(*u*) ≥ *ℓ*, so that *ℒ*_*ℓ*_ ⊆ *ℒ*_*ℓ*−1_. The *ℓ*th *contour* is the boundary of *ℒ*_*ℓ*_ (Fig. 3). Layer *ℒ*_*ℓ*_ (*ℓ* > 0) contains exactly those states that precede a match *m* with score *ℓ* ≤ *S*_*p*_(*m*) < *ℓ* + *r* (Lemma 5 in App. B.3).

#### Computing *S*_*p*_(*u*)

This last observation inspires our algorithm for computing chain scores. For each layer *ℒ*_*ℓ*_, we store the set *L*[*i*] of matches having score *ℓ*: *L*[*ℓ*] = {*m* ∈ ℳ ∣ *S*_*p*_(*m*) = *ℓ*}. The score *S*_*p*_(*u*) is then the highest *ℓ* such that layer *L*[*ℓ*] contains a match *m* reachable from *u* (*u* ⪯_*p*_ *m*). From Lemma 5 we know that *S*_*p*_(*u*) ≥ *ℓ* if and only if one of the layers *L*[*ℓ*^′^] for *ℓ*^′^ ∈ [*ℓ, ℓ* + *r*) contains a match preceded by *u*. We use this to compute *S*_*p*_(*u*) using a binary search over the layers *ℓ*. We initialize *L*[0]={*mω* } (*mω* is a fictive match at the target *vt*), sort all matches in ℳ by ⪯_*p*_, and process them in decreasing order (from the target to the start). After computing *S*_*p*_(end(*m*)), we add *m* to layer *S*_*p*_(*m*) = score(*m*) + *S*_*p*_(end(*m*)). Matches that do not precede the target (start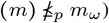 are ignored.

#### Pruning matches from *L*

When pruning matches starting or ending in state *u* in layer *ℓ*_*u*_ = *S*_*p*_(*u*), we remove all matches that start at *u* from layers *L*[*ℓ*_*u*_−*r*+1] to *L*[*ℓ*_*u*_], and all matches starting in some *v* and ending in *u* from layers *L*[*ℓ*_*v*_ −*r*+1] to *L*[*ℓ*_*v*_ ].

Pruning a match may change *Sp* in layers above *ℓu*, so we update them after each prune. We iterate over increasing *ℓ* starting at *ℓu* + 1 and recompute *ℓ*^′^ ≔ *S*_*p*_(*m*) ≤ *ℓ* for all matches *m* in *L*[*ℓ*]. If *ℓ*^′^ ≠ *ℓ*, we move *m* from *L*[*ℓ*] to *L*[*ℓ*^′^]. We stop iterating when either *r* consecutive layers were left unchanged, or when all matches in *r* − 1 + *ℓ* − *ℓ*^′^ consecutive layers have shifted down by the same amount *ℓ* − *ℓ*^′^. In the former case, no further scores can change, and in the latter case, *Sp* decreases by *ℓ* − *ℓ*^′^ for all matches with score ≥ *ℓ*. We remove the emptied layers *L*[*ℓ*^′^ + 1] to *L*[*ℓ*] so that all higher layers shift down by *ℓ* − *ℓ*^′^.

#### SH

Due to the simple structure of the seed heuristic, we also simplify its computation by only storing the start of each layer and the number of matches in each layer, as opposed to the full set of matches.

#### GCSH

Thm. 4 does not apply to gap-chaining seed heuristic since it uses chaining cost *γ*= max(*c*_gap_(*u, v*), *c*_seed_(*u, v*)) which is different from *c*_seed_(*u, v*). It turns out that in this new setting it is never optimal to chain two matches if the gap cost between them is higher than the seed cost. Intuitively, it is better to miss a match than to incur additional gapcost to include it. We capture this constraint by introducing a transformation *T* such that *u* ⪯_*T*_ *v* holds if and only if *c*_seed_(*u, v*) ≥ *c*_gap_(*u, v*), as shown in App. B.4. Using an additional *consistency* constraint on the set of matches we can compute 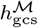 via *S*_*T*_ as before.

##### Definition 3

(Consistent matches). *A set of matches ℳ is* consistent *when for each m* ∈ ℳ *(from* ⟨*i, j*⟩ *to* ⟨*i*^′^, *j*^′^⟩*) with* score(*m*)>1, *for each* adjacent *pair of existing states* (⟨*i, j*±1⟩, ⟨*i*^′^, *j*^′^⟩) *and* (⟨*i, j*⟩, ⟨*i*^′^, *j*^′^±1⟩), *there is an* adjacent *match with corresponding start and end, and score at least* score(*m*)−1.

This condition means that for *r*=2, each exact match must be adjacent to four (or less around the edges of the graph) inexact matches starting or ending in the same state. Since we find all matches *m* with cm(*m*)<*r*, our initial set of matches is consistent. To preserve consistency, we do not prune matches if that would break the consistency of ℳ.

##### Definition 4

(Gap transformation). *The partial order* ⪯_*T*_ *on states is induced by comparing both coordinates after the* gap transformation

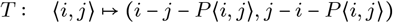

##### Theorem 5

*Given a* consistent *set of matches ℳ, the gap-chaining seed heuristic can be computed using scores in the transformed domain*:

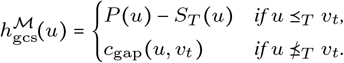

Using the transformation of the match coordinates, we can now reduce *c*_gs_ to *c*_seed_ and efficiently compute GCSH in any explored state.

## 4 Results

Our algorithm is implemented in the aligner A*PA^1^ in Rust. We compare it with state of the art exact aligners on synthetic (Sec. 4.2) and human (Sec. 4.3) data^2^ using PABENCH^3^. We justify our heuristics and optimizations by comparing their scaling and performance (Sec. 4.4).

### 4.1 Setup

#### Synthetic data

Our synthetic datasets are parameterized by sequence length *n*, induced error rate *e*, and total number of basepairs *N*, resulting in *N* /*n* sequence pairs. The first sequence in each pair is uniform-random from Σ^*n*^. The second is generated by sequentially applying ⌊*e*·*n*⌋ edit operations (insertions, deletions, and substitutions with equal 1/3 probability) to the first sequence. Introduced errors can cancel each other, making the *divergence d* between the sequences less than *e*. Induced error rates of 1%, 5%, 10%, and 15% correspond to divergences of 0.9%, 4.3%, 8.2%, and 11.7%, which we refer to as 1%, 4%, 8%, and 12%.

#### Human data

We use two datasets of ultra-long Oxford Nanopore Technologies (ONT) reads of the human genome: one without and one with genetic variation. All reads are 500–1100 kbp long, with mean divergence around 7%. The average length of the longest gap in the alignment is 0.1 kbp for ONT reads, and 2 kbp for ONT reads with genetic variation (detailed statistics in App. C.5). The reference genome is CHM13 (v1.1) (Nurk *et al*., 2022). The reads used for each dataset are:

- *ONT*: 50 reads sampled from those used to assemble CHM13.
- *ONT with genetic variation*: 48 reads from another human (Bowden *et al*., 2019), as used in the BIWFA paper (Marco-Sola *et al*., 2023).

#### Algorithms and aligners

We compare SH, CSH, and GCSH (all with pruning) as implemented in A*PA to the state-of-the-art exact aligners BIWFA and EDLIB. We also compare to Dijkstra’s algorithm and A* with previously introduced heuristics (gap cost and character frequencies of Hadlock (1988b), and SH without pruning of Ivanov *et al*. (2022)). We exclude SEQAN and PARASAIL since they are outperformed by WFA and EDLIB (Marco-Sola *et al*., 2021; Šošić and Šikić, 2017). We run all aligners with unit edit costs with traceback enabled.

#### A*PA parameters

Inexact matches (*r*=2) and short seeds (low *k*) increase the accuracy of GCSH for divergent sequences, thus reducing the number of expanded states. On the other hand, shorter seeds have more matches, slowing down precomputation and contour updates. A parameter grid search on synthetic data (App. C.1) shows that the runtime is generally insensitive to *k* as long as *k* is high enough to avoid too many spurious matches (*k* ≫ log_4_ *n*), and the potential is sufficiently larger than edit distance (*k* ≪ *r*/*d*). For *d*=4%, exact matches lead to faster runtimes, while *d*=12% requires *r*=2 and *k* < 2/*d* = 16.7. We fix *k* = 15 throughout the evaluations since this is a good choice for both synthetic and human data.

#### Execution

We use PABENCH on Arch Linux on an `Intel Core i7-10750H` processor with 64 GB of memory and 6 cores, without hyper-threading, frequency boost, and CPU power saving features. We fix the CPU frequency to `2.6GHz`, limit the memory usage to 32 GiB, and run 1 single-threaded job at a time with niceness −20.

#### Measurements

PaBench first reads the dataset from disk and then measures the wall-clock time and increase in memory usage of each aligner. Plots and tables refer to the average alignment time per aligned pair, and for A*PA include the time to build the heuristic. Best-fit polynomials are calculated via a linear fit in the log-log domain using the least squares method.

### 4.2 Scaling on synthetic data

#### Runtime scaling with length

We compare our A* heuristics with EDLIB, BIWFA, and other heuristics in terms of runtime scaling with *n* and *d* (Fig. 4, extended comparison in App. C.2). As theoretically predicted, EDLIB and BIWFA scale quadratically. For small edit distance, EDLIB is subquadratic due to the bit-parallel optimization. Dijkstra, A* with the gap heuristic, character frequency heuristic (Hadlock, 1988b), or original seed heuristic (Ivanov *et al*., 2022) all scale quadratically. The empirical scaling of A*PA is subquadratic for *d*≤12 and *n*≤10^7^, making it the fastest aligner for long sequences (*n*>30 kbp). For low divergence (*d*≤4%) even the simplest SH scales near-linearly with length (best fit *n*^1.06^ for *n*≤10^7^). For high divergence (*d*=12%) we need inexact matches, and the runtime of SH sharply degrades for long sequences (*n*>10^6^ bp) due to spurious matches. This is countered by chaining the matches in CSH and GCSH, which expand linearly many states (App. C.3). GCSH with DT is not exactly linear due to high memory usage and state reordering (App. C.7 shows the time spent on parts of the algorithm).

**Fig. 4.**
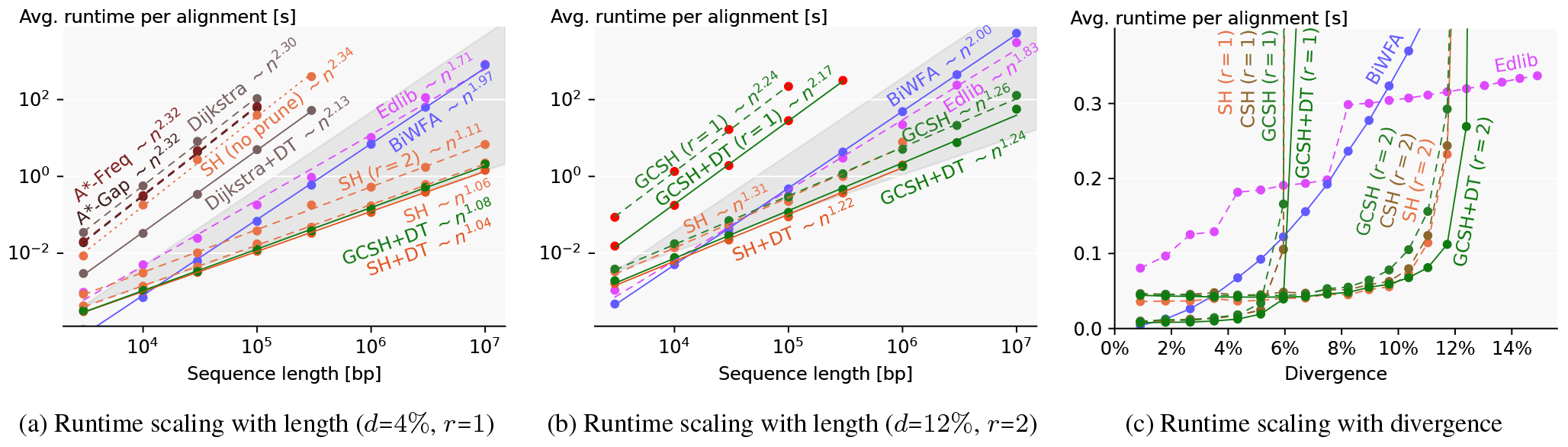
Runtime comparison on synthetic data. TODO LABELS (a)(b) Log-log plots comparing variants of our heuristic, including the simplest (SH) and most accurate (GCSH with DT), to EDLIB, BIWFA, and other algorithms (averaged over 10^6^ to 10^7^ total bp, seed length *k*=15). The slopes of the bottom (top) of the dark-grey cones correspond to linear (quadratic) growth. SH without pruning is dotted, and variants with DT are solid. For *d*=12%, red dots show where the heuristic potential is less than the edit distance. Missing data points are due to exceeding the 32 GiB memory limit. (c) Runtime scaling with divergence (*n*=10^5^, 10^6^ total bp, *k*=15).

#### Runtime scaling with divergence

Fig. 4c shows that A*PA has near constant runtime in *d* as long as the edit distance is sufficiently less than the heuristic potential (i.e. *d* ≪ *r*/*k*). In this regime, A*PA is faster than both EDLIB (linear in *d*) and BIWFA (quadratic in *d*). For 1 ≤ *d* ≤ 6%, exact matches have less overhead than inexact matches, while BIWFA is fastest for *d* ≤ 1%. A*PA becomes linear in *d* for *d* ≥ *r*/*k* (App. C.4).

#### Performance

A*PA with SH with DT is >500× faster than EDLIB and BIWFA for *d*=4% and *n*=10^7^ (Fig. 4a). For *n*=10^6^ and *d*≤12%, memory usage is less than 500 MB for all heuristics (App. C.6).

### 4.3 Speedup on human data

We compare runtime (Fig. 5, App. C.7), and memory usage (App. C.6) on human data. We configure A*PA to prune matches only when expanding their start (not their end), leaving some matches on the optimal path unpruned and speeding up contour updates. The runtime of A*PA (GCSH with DT) on ONT reads is less than EDLIB and BIWFA in all quartiles, with the median being >3× faster. However, the runtime of A*PA grows rapidly when *d*≥10%, so we set a time limit of 100 seconds per read, causing 6 alignments to time out. In real-world applications, the user would either only get results for a subset of alignments, or could use a different tool to align divergent sequences. With genetic variation, A*PA is 1.7× faster than EDLIB and BIWFA in median. Low-divergence alignments are faster than EDLIB, while high-divergence alignments are slower (3 sequences with *d*≥10% time out) because of expanding quadratically many states in complex regions (App. C.8). Since slow alignments dominate the total runtime, EDLIB has a lower mean runtime.

**Fig. 5.**
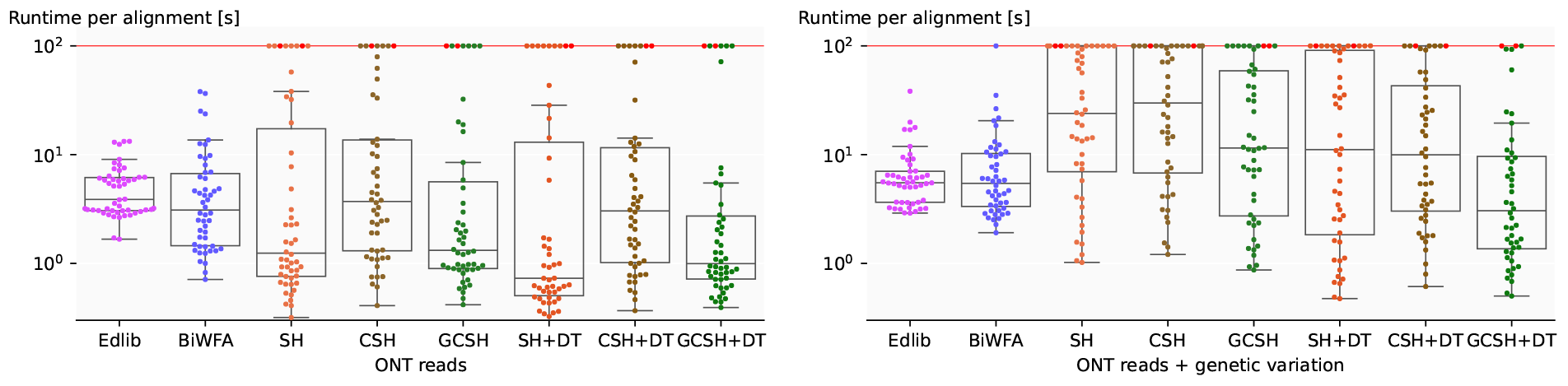
Runtime on long human reads. Each dot is an alignment without (left) or with (right) genetic variation. Runtime is capped at 100 s. Boxplots show the three quartiles, and red dots show where the edit distance is larger than the heuristic potential. The median runtime of A*PA (GCSH + DT, *k*=15, *r*=2) is 3× (left) and 1.7× (right) faster than EDLIB and BIWFA.

### 4.4 Effect of pruning, inexact matches, chaining, and DT

We visualize our techniques on a complex alignment in App. C.10.

#### SH with pruning enables near-linear runtime

Figure 4a shows that the addition of match pruning changes the quadratic runtime of SH without pruning to near-linear, giving multiple orders of magnitude speedup.

#### Inexact matches cope with higher divergence

Inexact matches double the heuristic potential, thereby almost doubling the divergence up to which A*PA is fast (Fig. 4c). This comes at the cost of a slower precomputation to find all matches.

#### Chaining copes with spurious matches

While CSH improves on SH for some very slow alignments (Fig. 5), more often the overhead of computing contours makes it slower than SH.

#### Gap-chaining copes with indels

GCSH is significantly and consistently faster than SH and CSH on human data, especially for slow alignments (Fig. 5). GCSH has less overhead over SH than CSH, due to filtering out matches 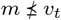.

#### Diagonal transition speeds up quadratic search

DT significantly reduces the number of expanded states when the A* search is quadratic (Fig. 4a and App. C.4). In particular, this results in a big speedup when aligning genetic variation containing big indels (Fig. 5).

CSH, GCSH, and DT only have a small impact on the uniform synthetic data, where usually either the SH is sufficiently accurate for the entire alignment and runtime is near-linear (*d* ≪ *r*/*k*), or even GCSH is not strong enough and runtime is quadratic (*d* ≫ *r*/*k*). On human data however, containing longer indels and small regions of quadratic search, the additional accuracy of GCSH and the reduced number of states explored by DT provide a significant speedup (App. C.10).

## 5 Discussion

### Seeds are necessary, matches are optional

The seed heuristic uses the lack of matches to penalize alignments. Given the admissibility of our heuristics, the more seeds without matches, the higher the penalty for alignments and the easier it is to dismiss suboptimal ones. In the extreme, not having any matches can be sufficient for finding an optimal alignment in linear time (App. C.9).

### Modes: Near-linear and quadratic

The A* algorithm with a seed heuristic has two modes of operation that we call *near-linear* and *quadratic*. In the near-linear mode A*PA expands few vertices because the heuristic successfully penalizes all edits between the sequences. When the divergence is larger than what the heuristic can handle, every edit that is not penalized by the heuristic increases the explored band, leading to a quadratic exploration similar to Dijkstra.

### Limitations

1. *Quadratic scaling*. Complex data can trigger a quadratic (Dijkstra-like) search, which nullifies the benefits of A* (Appendices C.8 and C.10). Regions with high divergence (*d*≤10%), such as high error rate or long indels, exceed the heuristic potential to direct the search and make the exploration quadratic. Low-complexity regions (e.g. with repeats) result in a quadratic number of matches which also take quadratic time.
2. *Computational overhead of A\**. Computing states sequentially (as in EDLIB, BIWFA) is orders of magnitude faster than computing them in random order (as in Dijkstra, A*). A*PA outperforms EDLIB and BIWFA (Fig. 4a) when the sequences are long enough for the linear-scaling to have an effect (*n*>30 kbp), and there are enough errors (*d*>1%) to trigger the quadratic behaviour of BIWFA.

### Future work

1. *Performance*. We are working on a DP-based version of A*PA that applies computational volumes (Spouge, 1989, 1991), block-based computations (Liu and Steinegger, 2023), and a SIMD version of EDLIB’s bit-parallelization (Myers, 1999). This has already shown 10× additional speedup on the human data sets and is less sensitive to the input parameters. Independently, the number of matches could be reduced by using variable seed lengths and skipping seeds with many matches.
2. *Generalizations*. Our chaining seed heuristic could be generalized to non-unit and affine costs, and to semi-global alignment. Cost models that better correspond to the data can speed up the alignment.
3. *Relaxations*. At the expense of optimality guarantees, inadmissible heuristics could speed up A* considerably. Another possible relaxation would be to validate the optimality of a given alignment instead of computing it from scratch.
4. *Analysis*. The near-linear scaling with length of A* is not asymptotic and requires a more thorough theoretical analysis (Medvedev, 2022a).

## Acknowledgements

We are grateful to Mykola Akulov for his help with Figs. 1 and 3, to Daniel Liu for his involvement in developing PABENCH, to Benjamin Bichsel, Maximilian Mordig, André Kahles, and the anonymous reviewers for valuable comments on drafts, and to Sergey Nurk for his help with the human data. R. Groot Koerkamp is financed by ETH Research Grant ETH-17 21-1 to Gunnar Rätsch. *Conflict of Interest:* none declared.

## A Pseudocode

### A.1 A* algorithm for match pruning

We present our variant of A* (Hart *et al*., 1968) that supports match pruning (Algorithm 1). All computed values of *g* are stored in a hash map, and all *open* states are stored in a bucket queue of tuples (*v, g*(*v*), *f* (*v*)) ordered by increasing *f*. Line 14 prunes (removes) a match and thereby increases some heuristic values before that match. As a result, some *f* -values in the priority queue may become outdated. Line 11 solves this problem by checking if the *f* -value of the state about to be expanded was changed, and if so, line 12 pushes the updated state to the queue, and proceeds by choosing a next best state. This way, we guarantee that the expanded state has minimal updated *f*. To reconstruct the best alignment we traceback from the target state using the hash map *g* (not shown).

### A.2 Diagonal transition for A*

For a given distance *g*, the diagonal transition method only considers the *farthest-reaching* (f.r.) state *u*=⟨*i, j*⟩ on each diagonal *k*=*i*−*j* at distance *g*. We use *F*_*gk*_ ≔ *i*+*j* to indicate the antidiagonal^4^ of the farthest reaching state. Let *X*_*gk*_ be the farthest state on diagonal *k* adjacent to a state at distance *g*−1, which is then *extended* into *F*_*gk*_ by following as many matches as possible. The edit distance is the lowest *g* such that *F*_*g,n*−*m* ≥ *n* + *m*_, and we have the recursion

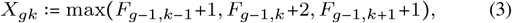

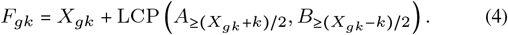

The base case is *X*_0,0_=0 with default value *F*_*gk*_=−∞ for *k*>∣*g*∣, and LCP is the length of the longest common prefix of two strings. Each edge in a traceback path is either a match created by an extension (4), or a mismatch starting in a f.r. state (3). We call such a path an *f*.*r. path*.

#### Algorithm 1

A* algorithm with match pruning.

Lines added for pruning (11, 12, and 14) are marked in **bold**.

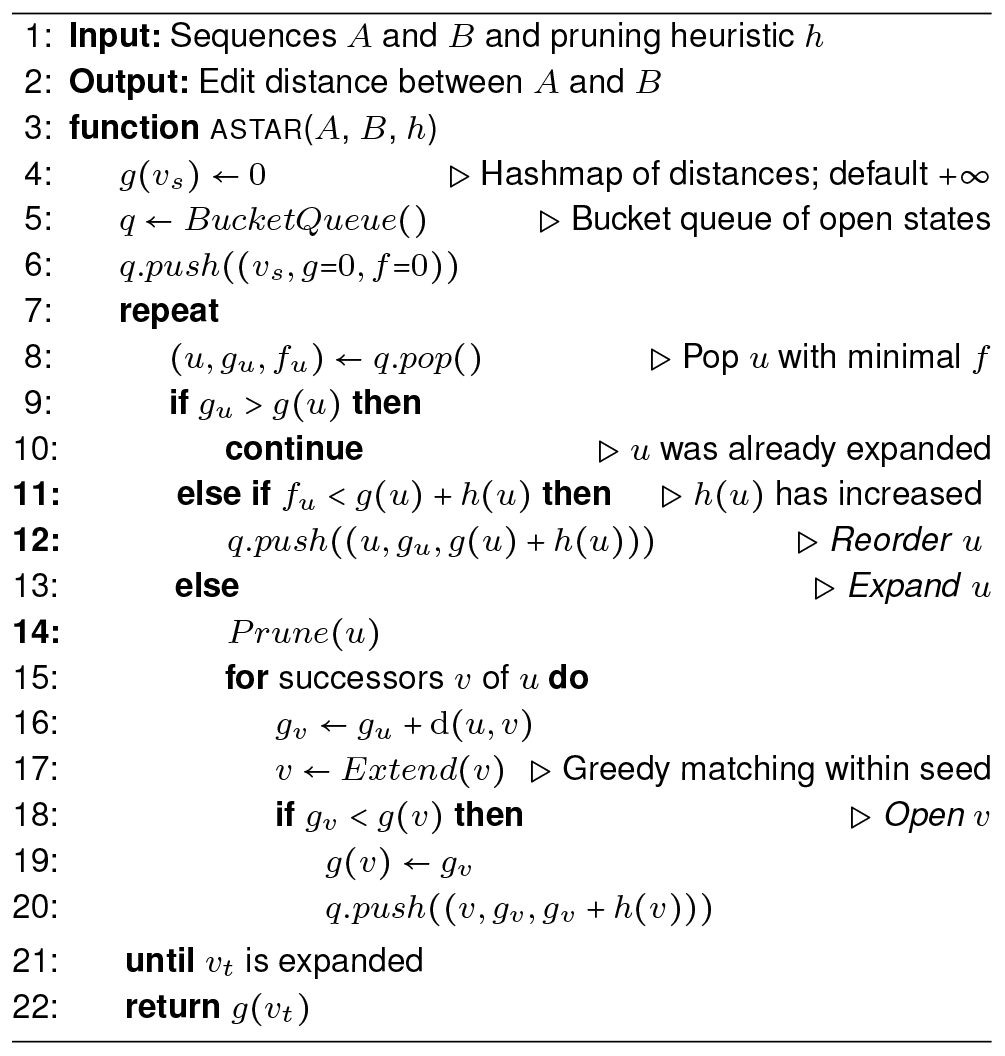

#### Implementation

In Algorithm 2 we further modify the A* algorithm to only consider f.r. paths. We replace the map *g* that tracks the best distance by a map *F*_*gk*_ that tracks f.r. states (lines 4, 18, and 19). Instead of *g*(*u*) decreasing over time, we now ensure that *F*_*g,k*_ increases over time. Each time a state *u* is opened or expanded, the check whether *g*(*u*) decreases is replaced by a check whether *F*_*gk*_ increases (line 9). This causes the search to skip states that are known to not be farthest reaching. The proof of correctness (Thm. 2) still applies.

Alternatively, it is also possible to implement A* directly in the diagonal-transition state-space by pushing states *F*_*gk*_ to the priority queue, but for simplicity we keep the similarity with the original A*.

### A.3 Implementation notes

Here we list implementation details on performance and correctness.

#### Bucket queue

We use a hashmap to store all computed values of *g* in the A* algorithm. Since the edit costs are bounded integers, we implement the priority queue using a *bucket queue* (Hitchner, 1968; Dial, 1969; Bertsekas, 1991). Unlike heaps, this data structure has amortized constant time push and pop operations since the value difference between consecutive pop operations is bounded.

#### Greedy matching of letters

From a state ⟨*i, j*⟩ where *a*_*i*_ = *b*_*j*_, it is sufficient to only consider the matching edge to ⟨*i*+1, *j*+1⟩ (Allison, 1992; Ivanov *et al*., 2020), and ignore the insertion and deletion edges to ⟨*i, j*+1⟩ and ⟨*i*+1, *j*⟩. During alignment, we greedily match as many letters as possible within the current seed before inserting only the last open state in the priority queue, but we do not cross seed boundaries in order to not interfere with match pruning. This optimization is superseded by the DT-optimization. We include greedily matched states in the reported number of expanded states.

#### Priority queue offset

Pruning the last remaining match in a layer may cause an increase of the heuristic in all states preceding the start *u* of the match. This invalidates *f* values in the priority queue and causes reordering. We skip most of the update operations by storing a global offset to the *f* -values in the priority queue, which we update when all states in the priority queue precede *u*.

##### Algorithm 2

A*-DT algorithm with match pruning.

Lines changed for diagonal transition (4, 9, 18, and 19) are in **bold**.

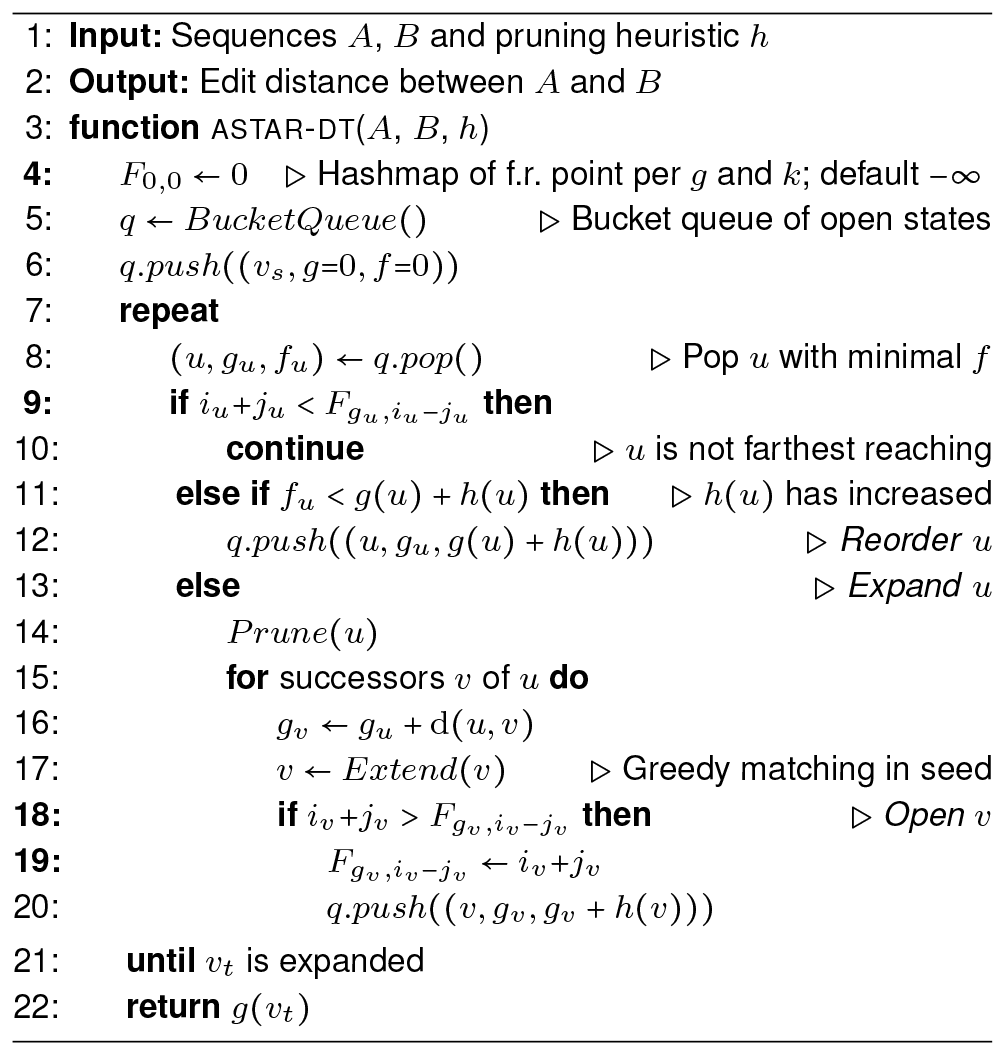

#### Split vector for layers

Pruning a match may trigger the removal of one or more layers of matches in *L*, and the shifting down of higher layers. To efficiently remove layers, we use a *split vector* data structure consisting of two stacks. In the first stack we store the layers before the last deleted layer, and in the second stack the remaining layers in reverse order. Before deleting a layer, we move layers from the top of one of the stacks to the top of the other, until the layer to be deleted is at the top of one of the stacks. Removing layers in decreasing order of *ℓ* takes linear total time.

#### Binary search speedup

Instead of using binary search to determine the layer/score *S*_*p*_(*u*) (Algorithm 3), we first try a linear search around either the score of the parent of *u* or a known previous score at *u*. This linear search usually finds the correct layer in a few iterations, or otherwise we fall back to binary search.

In practice, most pruning happens near the tip of the search, and the number of layers between the start *v*_*s*_ and an open state *u* rarely changes. Thus, to account for changing scores, we store a *hint* of value *S*_*p*_(*v*_*s*_) − *S*_*p*_(*u*) in a hashmap and start the search at *S*_*p*_(*v*_*s*_) − *hint*.

#### Code correctness

Our implementation A*PA is written in Rust, contains many assertions testing e.g. the correctness of our A* and layers data structure implementation, and is tested for correctness and performance on randomly-generated sequences. Correctness is checked against simpler algorithms (Needleman-Wunsch) and other aligners (EDLIB, BIWFA).

### A.4 Computation of the heuristic

Algorithm 3 shows how the heuristic is initialized, how *S*_*p*_(*u*) and *h*(*u*) are computed, and how matches are pruned.

### A.5 Worst-case runtime asymptotics

Our algorithms optimize for the average case performance. Nevertheless, we discuss the worst-case time of each part of the algorithm. To analyze the worst-case asymptotics of A*PA, let *M* = *O*(*n*^2^) be the number of (inexact) matches, and let *E* = *O*(*n*^2^) be the number of expanded states. Then the asymptotical runtime of our algorithm breaks down as:

1. Finding all matches takes *O*(*n* + *M*) time.
2. Building the contours datastructure takes *O*(*M* log *M*) time.
3. Each expanded state is pushed onto and popped from the priority queue (*O*(1) using a bucket queue), looked up in a hashmap (*O*(1)), and has its neighbours explored (*O*(1)), for *O*(*E*) total.
4. Evaluating the heuristic requires a binary search over contours, which is bounded by *O*(*n* lg *n*), since we test at most lg *n* contours each containing at most *n* matches. (This could be improved to *O*(log^2^ *n*) by storing each contour as an ordered set, but it is not usually a bottleneck.)
5. Pruning a match requires a dictionary lookup and change for *O*(1). Updating contours after pruning requires iterating over higher contours until no more changes are triggered. This could take *O*(*M*) for each match that is pruned, leading to a naive upper bound of *O*(*M* ^2^).
6. Reordering states takes *O*(*En*), since the heuristic in each expanded state can decrease at most *n* times.
7. The traceback requires *O*(*n*) time.

Taking this together, we obtain *O*(*n*+*M* log *M* +*En* lg *n*+*M* ^2^+*En*). When the number of matches *M* is Θ(*n*^2^), the updating of contours has the worst upper bound and the asymptotics becomes *O*(*n*^4^). Whether this upper bound can be reached in practice or whether a tigher bound exists needs further investigation. Even if the worst-case was Θ(*n*^2^), in practice our algorithm only runs efficiently in regimes well below this bound.

App. C.1 discusses the effect of *r* and *k* on the number of expanded states when aligning synthetic data, and App. C.4 discusses the effect of the divergence *d* on the number of expanded states.

#### Algorithm 3

Computation of the heuristic.

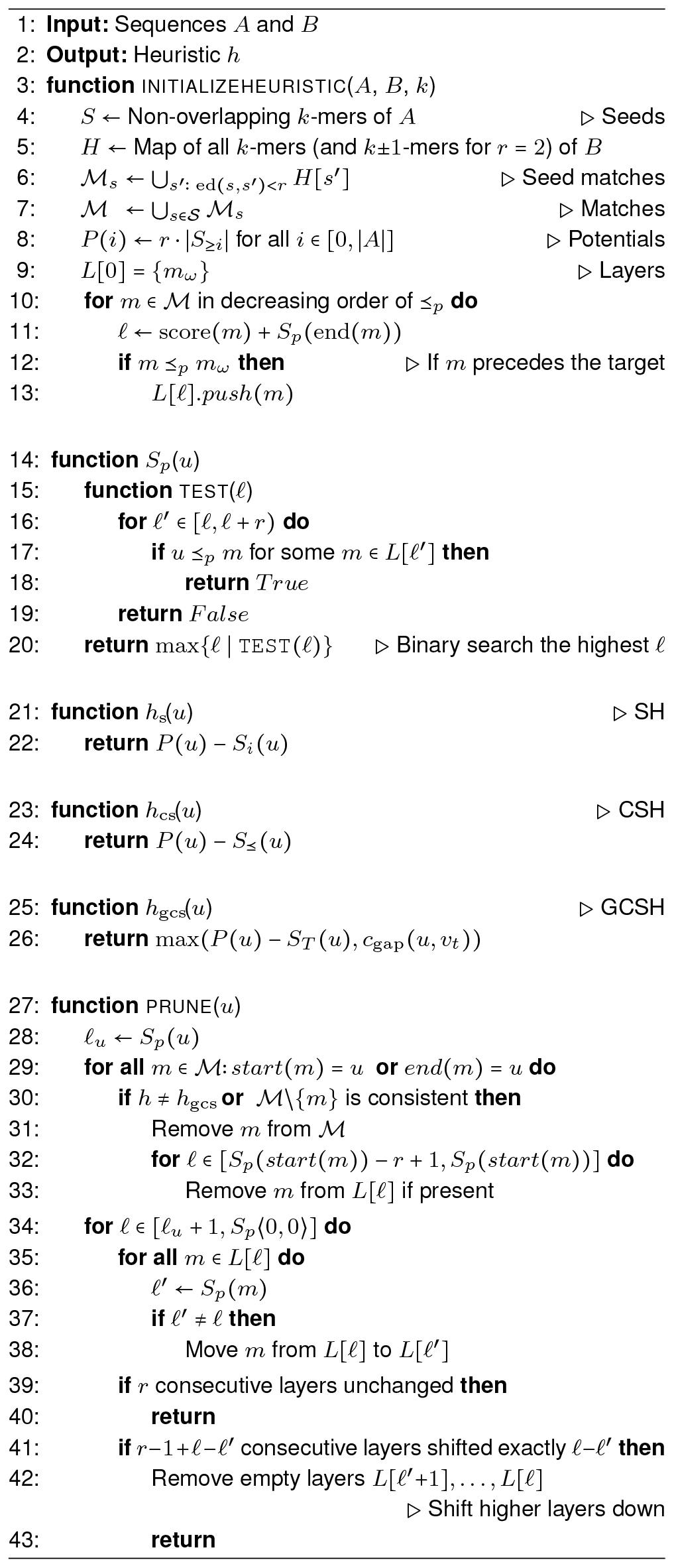

## B Proofs

### B.1 Admissibility

Our heuristics are not *consistent*, but we show that a weaker variant holds for states *at the start of a seed*.

#### Definition 5

(Start of seed). *A state* ⟨*i, j*⟩ *is* at the start of some seed *when i is a multiple of the seed length k, or when i* = *n*.

#### Lemma 1

(Weak triangle inequality). *For states u, v, and w with v and w at the starts of some seeds, all γ* ∈ {*c*_seed_, *c*_gap_, *c*_gs_} *satisfy*

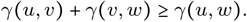

*Proof*. Both *v* and *w* are at the start of some seeds, so for *γ* = *c*_seed_ we have the equality *c*_seed_(*u, w*) = *c*_seed_(*u, v*) + *c*_seed_(*v, w*).

For *γ* = *c*_gap_,

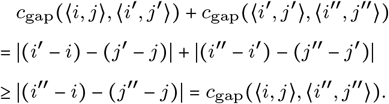

And lastly, for *γ* = *c*_gs_,

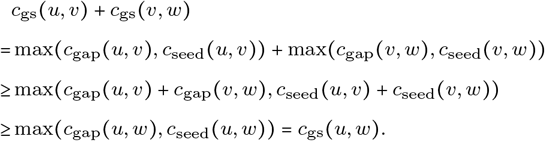

#### Lemma 2

(Weak consistency). *Let* 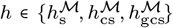 *be a heuristic with partial order* ⪯_*p*_, *and let u* ⪯_*p*_ *v be states with v at the start of a seed. When there is a shortest path π*^∗^ *from u to v such that ℳ contains all matches of cost less than r on π*^∗^, *it holds that h*(*u*) ≤ d(*u, v*)+*h*(*v*).

*Proof*. The path *π*^∗^ covers each seed in 𝒮*u*…*v* that must to be fully aligned between *u* and *v*. Since the seeds do not overlap, their shortest alignments 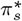 in *π*^∗^ do not have overlapping edges. Let *u* ⪯ *m*_1_ ⪯_*p*_ ⋅⋅⋅⪯_*p*_ *m*_*l*_ ⪯_*p*_ *v* be the chain of matches *m*_*i*_ ∈ ℳ corresponding to those 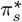 of cost less than *r* (Fig. 6). Since the matches and the paths between them are disjoint, c_path_(*π*^∗^) is at least the cost of the matches c_m_(*m*_*i*+1_) = d(start(*m*_*i*+1_), end(*m*_*i*+1_)) plus the cost to chain these matches *γ*(end(*m*_*i*_), start(*m*_*i*+1_)) ≤ d(end(*m*_*i*_), start(*m*_*i*+1_)). Putting this together:

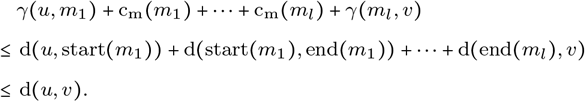

Now let *v* ⪯_*p*_ *m*_*l*+1_ ⪯_*p*_ ⋅⋅⋅⪯_*p*_ *ml*′ ⪯_*p*_ *v*_*t*_ be a chain of matches minimizing *h*(*v*) (Def. 1) with *w* ≔ start(*m*_*l*+1_). This chain also minimizes *h*(*w*) and thus *h*(*v*) = *γ*(*v, w*) + *h*(*w*). We can now bound the cost of the joined chain from *u* to *v* and from *w* to the end and get our result via *γ*(*m*_*l*_, *w*) ≤ *γ*(*m*_*l*_, *v*) + *γ*(*v, w*) (Lemma 1)

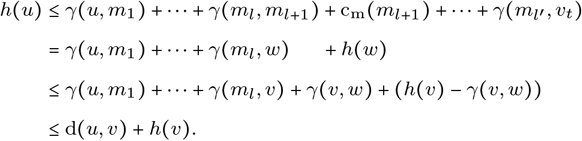

#### Theorem 1

*The seed heuristic h_s_, the chaining seed heuristic h_cs_, and the gap-chaining seed heuristic h_gcs_ are admissible. Furthermore, 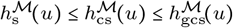 for all states u.*

*Proof*. We will prove 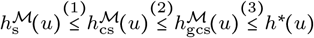, which implies the admissibility of all three heuristics.

1. Note that *u* ⪯ *v* implies *u* ⪯_*i*_ *v* and hence any ⪯-chain is also a ⪯_*i*_-chain. A minimum over the superset of ⪯_*i*_-chains is at most the minimum of the subset of ⪯-chains, and hence 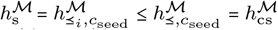.
2. The only difference between 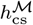 and 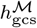 is that the former uses cs gcs *c*_seed_ and the latter uses the gap-seed cost *c*_gs_ ≔ max(*c*_gap_, *c*_seed_). Since *c*_seed_ ≤ *c*_gs_ we have 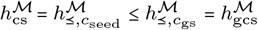.
3. When ℳ is the set of all matches with costs strictly less than *r*, admissibility follows directly from Lemma 2 with *v* = *vt* via

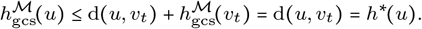

**Fig. 6.**
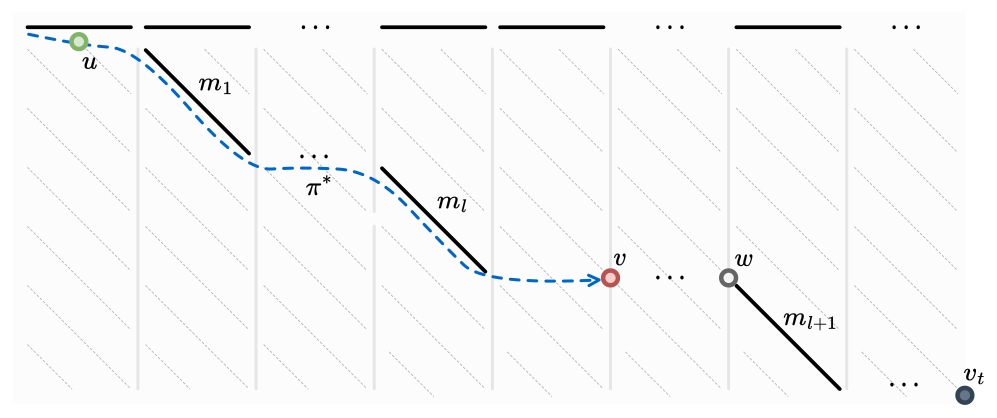
Variables of the proof of Lemma 2.

### B.2 Match pruning

During the A* search, we continuously improve our heuristic using match pruning. The pruning increases the value of our heuristics and breaks their admissibility. Nevertheless, we prove in two steps that A* with match pruning still finds a shortest path. First, we introduce the concept of a *weakly-admissible heuristic* and show that A* using a weakly-admissible heuristic finds a shortest path (Thm. 2). Then, we show that our pruning heuristics are indeed weakly admissible (Thm. 3).

**A* with a weakly-admissible heuristic finds a shortest path**.

#### Definition 6

(Fixed vertex). *A vertex u is* fixed *if it is expanded and A* has found a shortest path to it, that is, g*(*u*) = *g*^∗^(*u*).

A fixed vertex cannot be opened again (Algorithm 1, line 18), and hence remains fixed.

#### Definition 7

(Weakly admissible). *A heuristic ĥ is* weakly admissible *if at any moment during the A* search there exists a shortest path π*^∗^ *from vs to vt in which all vertices u* ∈ *π*^∗^ *after its last fixed vertex n*^∗^ *satisfy ĥ*(*u*) ≤ *h*^∗^(*u*).

To prove that A* finds a shortest path when used with a weakly-admissible heuristic, we follow the structure of Hart *et al*. (1968). First we restate their Lemma 1 in our notation with a slightly stronger conclusion that follows directly from their proof.

#### Lemma 3

(Lemma 1 of Hart *et al*. (1968)). *For any unfixed vertex n and for any shortest path π*^∗^ *from vs to n, there exists an open vertex n*^′^ *on π*^∗^ *with g*(*n*^′^) = *g*^∗^(*n*^′^) *such that π*^∗^ *does not contain fixed vertices after n*^′^.

Next, we prove that in each step the A* algorithm can proceed along a shortest path to the target:

#### Corollary 1

(Generalization of Corollary to Lemma 1 of Hart *et al*. (1968)). *Suppose that ĥis weakly admissible, and that A* has not terminated. Then, there exists an open vertex n*^′^ *on a shortest path from v*_*s*_ *to v*_*t*_ *with f* (*n*^′^) ≤ *g*^∗^(*vt*).

*Proof*. Let *π*^∗^ be the shortest path from *vs* to *vt* given by the weak admissibility of *ĥ*(Def. 7). Since A* has not terminated, *vt* is not fixed. Substitute *n* = *vt* in Lemma 3 to derive that there exists an open vertex *n*^′^ on *π*^∗^ with *g*(*n*^′^) = *g*^∗^(*n*^′^). By definition of *f* we have *f* (*n*^′^) = *g*(*n*^′^) + ĥ(*n*^′^). Since *π*^∗^ does not contain any fixed vertices after *n*^′^, the weak admissibility of *ĥ*implies *ĥ*(*n*^′^) ≤ *h*^∗^(*n*^′^). Thus, *f* (*n*^′^) = *g*(*n*^′^) + ĥ(*n*^′^) ≤ *g*^∗^(*n*^′^) + *h*^∗^(*n*^′^) = *g*^∗^(*v*_*t*_).

#### Theorem 2

*A* with a weakly-admissible heuristic finds a shortest path*.

*Proof*. The proof of Theorem 1 in Hart *et al*. (1968) applies, with the remark that instead of an arbitrary shortest path, we use the specific path *π*^∗^ given by the weak admissibility and the specific vertex *n*^′^ given by Corollary 1.

#### Our heuristics are weakly admissible

A *consistent* heuristic finds the correct distance to each vertex as soon as it is expanded. While our heuristics are not consistent, this property is true for states *at the starts of seeds* (when *i* is a multiple of the seed length *k*, or when *i* = *n*).

##### Lemma 4

*For ĥ* ∈ {ĥ_s_, *ĥ*_cs_, *ĥ*_gcs_}, *every state at the start of a seed becomes fixed immediately when A* expands it*.

*Proof*. We use a proof by contradiction: suppose that *v* is a state at the start of some seed that is expanded but not fixed. In other words, *f* (*v*) is minimal among all open states, but the shortest path *π*^∗^ from *vs* to *v* has strictly smaller length *g*^∗^(*v*) < *g*(*v*).

Let *n*^∗^ be the last fixed state on *π*^∗^ before *v*, and let *u* ∈ *π*^∗^ be the successor of *n*^∗^. State *u* is open because its predecessor *n*^∗^ is fixed and on a shortest path to *u*. Let the chain of all matches of cost less than *r* on *π*^∗^ between *u* and *v* be *u* ⪯ *m*_1_ ⪯ ⋅⋅⋅⪯ *m*_*l*_ ⪯ *v*. Since *n*^∗^ is the last fixed state on *π*^∗^, none of these matches has been pruned, and they are all in ĥ/*E* as well. This means we can apply Lemma 2 to get *h*(*u*) ≤ d(*u, v*)+*h*(*v*), so

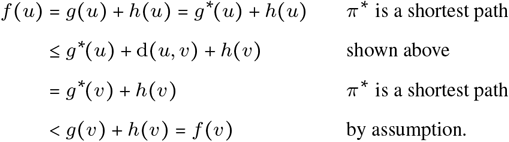

This proves that *f* (*u*) < *f* (*v*), resulting in a contradiction with the assumption that *v* is an open state with minimal *f*.

##### Theorem 3

*The pruning heuristics ĥ*_s_, *ĥ*_cs_, *ĥ*_gcs_ *are weakly admissible*.

*Proof*. Let *π*^∗^ be a shortest path from *vs* to *vt*. At any point during the A* search, let *n*^∗^ be the farthest expanded state on *π*^∗^ that is at the start of a seed. By Lemma 4, *n*^∗^ is fixed. By the choice of *n*^∗^, no states on *π*^∗^ after *n*^∗^ that are at the start of a seed are expanded, so no matches on *π*^∗^ following *n*^∗^ are pruned. Now the proof of Thm. 1 applies to the part of *π*^∗^ after *n*^∗^ without changes, implying that *ĥ*(*u*) ≤ *h*^∗^(*u*) for all *u* on *π*^∗^ following *n*^∗^, for any *ĥ* ∈ *ĥ*_s_, *ĥ*_cs_, *ĥ*_gcs_ }.

### B.3 Computation of the (chaining) seed heuristic

#### Theorem 4

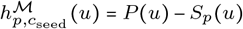 *for any partial order* ⪯_*p*_ *that is a refinement of* ⪯_*i*_ *(i*.*e. u* ⪯_*p*_ *v must imply u* ⪯_*i*_ *v)*.

*Proof*. For a chain of matches {*m*_*i*_} ⊆ ℳ, let *s*_*i*_ and *t*_*i*_ be the start and end states of *m*_*i*_. We translate the terms of our heuristic from costs to potentials and match scores (Sec. 3.3):

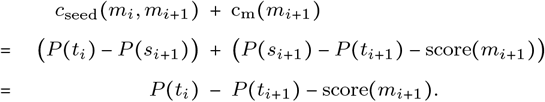

The heuristic (Def. 1) can then be rewritten as follows:

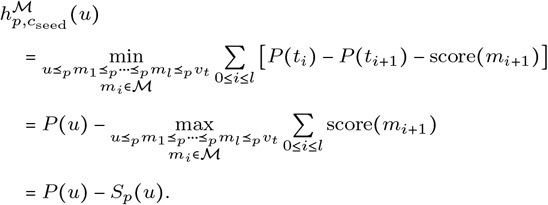

#### Lemma 5

*Layer ℒ*_*ℓ*_ *(ℓ* > 0*) is fully determined by the set of those matches m for which ℓ* ≤ *Sp*(*m*) < *ℓ* + *r:*

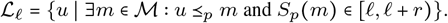

*Proof*. Take any state *u* ∈ *ℒ*_*ℓ*_. Its score *S*_*p*_(*u*) ≥ *ℓ* > 0 implies that there is a non-empty ⪯_*p*_-chain *u* ⪯_*p*_ *m*_1_ ⪯_*p*_ *m*_2_ ⪯_*p*_ … with score(*m*_1_)+ score(*m*_2_) +⋅⋅⋅≥ *ℓ*. The score of each match is less than *r* and thus there must be a match *m*_*i*_ so that the subset of the chain starting at *m*_*i*_ has score *S*_*p*_(*m*_*i*_) = score(*m*_*i*_) + score(*m*_*i*+1_) + ⋅⋅⋅+ in the interval [*ℓ, ℓ* + *r*). This implies that for any *u* with score *Sp*(*u*) ≥ *ℓ* > 0 there is a match with score in [*ℓ, ℓ* + *r*) succeeding *u*, as required.

### B.4 Computation of the gap-chaining seed heuristic

In this section we prove (Thm. 5) that we can change the dependency of GCSH on *c*_gs_ to *c*_seed_ by introducing a new partial order ⪯_*T*_ on the matches. This way, Thm. 4 applies and we can efficiently compute GCSH. Recall that the gap-seed cost is *c*_gs_ = max(*c*_gap_, *c*_seed_), and that the gap transformation is:

#### Definition 4

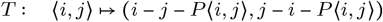

The following lemma allows us to determine whether *c*_gap_ or *c*_seed_ dominates the cost *c*_gs_ between two matches, based on the relation ⪯_*T*_.

#### Lemma 6

*Let u and v be two states with v at the start of some seed. Then u* ⪯_*T*_ *v if and only if c*_gap_(*u, v*) ≤ *c*_seed_(*u, v*). *Furthermore, u* ⪯_*T*_ *v implies u* ⪯ *v*.

*Proof*. Let *u* = ⟨*i, j*⟩ and *v* = ⟨*i*^′^, *j*^′^⟩. By definition, *u* ⪯_*T*_ *v* is equivalent to

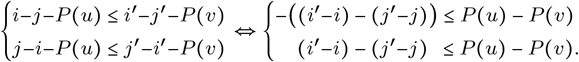

This simplifies to

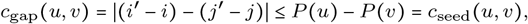

where the last equality holds because *v* is at the start of a seed.

For the second part, *u* ⪯_*T*_ *v* implies 0 ≤ *c*_gap_(*u, v*) ≤ *c*_seed_(*u, v*) = *P* (*u*) − *P* (*v*) and hence *P* (*u*) ≥ *P* (*v*). Since *v* is at the start of a seed, this directly implies *i* ≤ *i*^′^. Since seeds have length *k* ≥ *r* we have

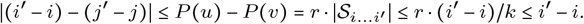

This implies *j*^′^ − *j* ≥ 0 and hence *j* ≤ *j*^′^, as required.

A direct corollary is that for *u* ⪯ *v* with *v* at the start of some seed, we have

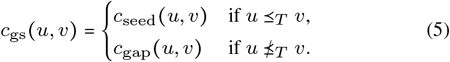

A second corollary is that start(*m*) ⪯_*T*_ end(*m*) for all matches *m*∈ℳ, since a match from *u* to *v* satisfies *c*_gap_(*u, v*) < *r* = *c*_seed_(*u, v*) by definition.

#### Lemma 7

*When the set of matches* ℳ *is consistent*, 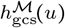*can be computed using* ⪯_*T*_ *-chains only:*

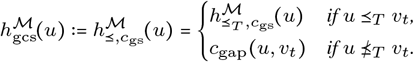

*Proof*. We write 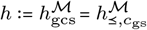 (Def. 1) and 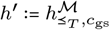

Case 1: 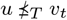. Let *u* ⪯ *m*_1_ ⪯ ⋅⋅⋅⪯ *m*_*l*_ ⪯ *v*_*t*_ be a chain minimizing *h*(*u*) in Def. 1, so

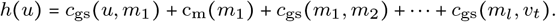

By definition, *c*_gs_ ≥ *c*_gap_, and c_m_(*m*_*i*_) ≥ *c*_gap_(start(*m*_*i*_), end(*m*_*i*_)), so the weak triangle inequality (Lemma 1) for *c*gap gives

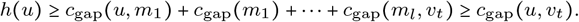

Since 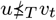, the empty chain *u*⪯*v*_*t*_ has cost *h*(*u*) ≤ *c*_gs_(*u, v*_*t*_) = *c*gap(*u, vt*). Combining the two inequalities, *h*(*u*) = *c*gap(*u, vt*).

Case 2: *u* ⪯_*T*_ *v*_*t*_. First rewrite *h* and *h*^′^ recursively as

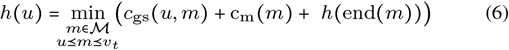

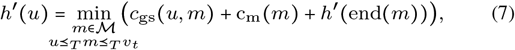

both with base case *h*(*vt*)=*h*^′^(*vt*) = 0 after eventually taking *mω*. We will show that

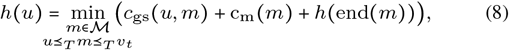

which is exactly the recursion for *h*^′^, so that by induction *h*(*u*) = *h*^′^(*u*). By Lemma 6, every ⪯_*T*_ -chain is a ⪯-chain, so *h*(*u*) ≤ *h*^′^(*u*). To prove *h*(*u*)=*h*^′^(*u*), it remains to show the reverse inequality, *h*^′^(*u*) ≤ *h*(*u*). To this end, choose a match *m* that

(priority 0) minimizes *h*(*u*) in Eq. (6), and among those, has (priority 1) maximal *c*_seed_(*u, m*), and among those, has (priority 2) minimal *c*gap(*u, m*), and among those, has (priority 3) minimal *c*gap(*m, vt*).

We show that *u* ⪯_*T*_ *m* (in 2.A) and *m* ⪯_*T*_ *v*_*t*_ (in 2.B), which proves Eq. (8).

Part 2.A: *u* ⪯_*T*_ *m*. Let *s* and *t* be the begin and end of *m* (Fig. 7), and let *m*^′^ be a match minimizing *h*(*t*) in Eq. (6) so

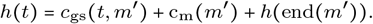

**Fig. 7.**
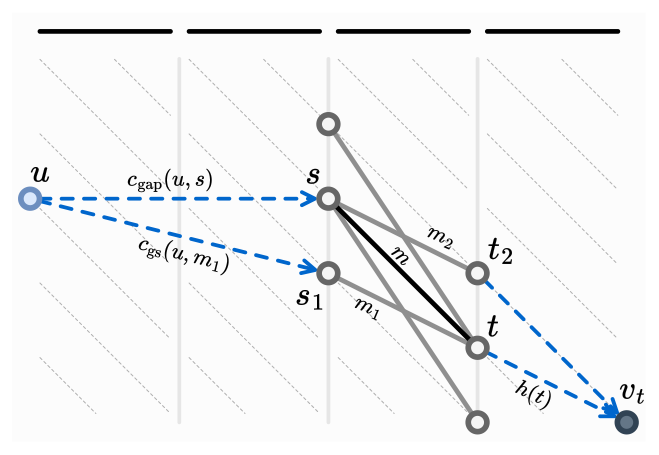
Variables in Case 2 of the proof of Lemma 7. Match *m* has score(*m*) = 2, so it has adjacent matches *m*1 and *m*2 (gray) with score(*mi*) ≥ 1.

Since *m*^′^ comes after *t* we have *c*_seed_(*u, m*^′^) > *c*_seed_(*u, m*) (p.1) and hence *m*^′^ does not minimize *h*(*u*) (p.0):

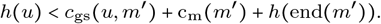

Using the minimality of *m*, the non-minimality of *m*^′^, and the triangle inequality we get

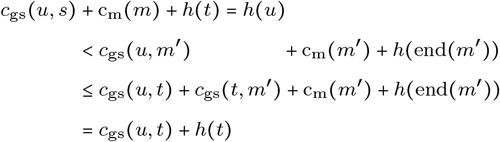

so we have

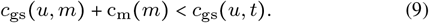

From the triangle inequality for *c*_gap_, from *c*_gap_ (*u, s*) ≤ *c*_gs_(*u, s*), and from *c*_gap_(*s, t*) ≤ c_m_(*m*), and from Eq. (9) we obtain

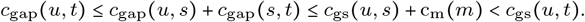

This implies *c*_gs_(*u, t*) = *c*_seed_(*u, t*) and hence reusing Eq. (9)

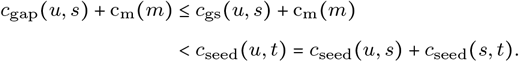

We have c_m_(*m*) = *c*_seed_(*s, t*) − score(*m*), so the above simplifies to *c*_gap_(*u, s*) < *c*_seed_(*u, s*) + score(*m*) and since these are integers *c*_gap_(*u, s*) ≤ *c*_seed_(*u, s*) + score(*m*) − 1.

When score(*m*) = 1, this implies *c*_gap_(*u, s*) ≤ *c*_seed_(*u, s*) and *u* ⪯_*T*_ *s* = start(*m*) by Lemma 6.

When score(*m*) > 1, suppose that *c*_gap_(*u, s*) > *c*_seed_(*u, s*) ≥ 0. That means that *u* is either above or below the diagonal of *s*. Let *s*_1_ = ⟨*s*_*i*_, *s*_*j*_ ± 1⟩ be the state adjacent to *s* on the same side of this diagonal as *u*. This state exists since *u* ⪯ *s*_1_ ⪯ *t*. Then *c*_gap_(*u, s*_1_) = *c*_gap_(*u, s*)−1, and by consistency of ℳ there is a match *m*_1_ from *s*_1_ to *t* with c_m_(*m*_1_) ≤ c_m_(*m*)+1. Then

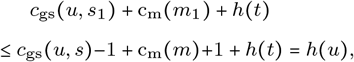

showing that *m*_1_ minimizes *h*(*u*) (p.0). Also *c*_seed_(*u, m*_1_) = *c*_seed_(*u, m*_1_) (p.1) and *c*_gap_(*u, m*_1_) < *c*_gap_(*u, m*) (p.2), so that *m*_1_ contradicts the minimality of *m*. Thus, *c*_gap_(*u, s*) > *c*_seed_(*u, s*) is impossible and *u* ⪯_*T*_ *s*.

Part 2.B: *m* ⪯_*T*_ *v*_*t*_. When there is some match *m*^′^ succeeding *m* in the chain, we have *m* ⪯ *m*^′^ ⪯ *vt* and hence *m* ⪯ *vt*. Thus, suppose that *m* is the only match in the chain *u* ⪯ *m* ⪯ *vt* minimizing *h*(*u*). We repeat the proofs of Part 2.A in the reverse direction to show that *m* ⪯_*T*_ *v*_*t*_.

Since *c*_seed_(*u, m*) is maximal, *h*(*u*) < *c*_gs_(*u, m*_*ω*_) and thus

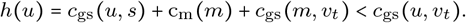

By assumption we have *u* ⪯_*T*_ *v*_*t*_ and thus *c*_gs_(*u, v*_*t*_) = *c*_seed_(*u, v*_*t*_). This gives

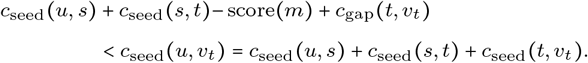

Cancelling terms we obtain *c*_gap_(*t, v*_*t*_) < *c*_seed_(*t, v*_*t*_) + 1 and since they are integers *c*_gap_(*t, v*_*t*_) ≤ *c*_seed_(*t, v*_*t*_) + score(*m*)−1. When score(*m*) = 1, this implies *t* ⪯_*T*_ *v*_*t*_, as required.

When score(*m*) > 1, suppose that *c*_gap_(*t, v*_*t*_) > *c*_seed_(*t, v*_*t*_). Let *t*_2_ = ⟨*t*_*i*_, *t*_*j*_ ± 1⟩ be the state adjacent to *t* on the same side of the diagonal as *v*_*t*_. By consistency of M there is a match *m*_2_ from *s* to *t*_2_ with score(*m*_2_) ≥ score(*m*)−1 and *c*_gap_(*t*_2_, *v*_*t*_) = *c*_gap_(*t, v*_*t*_) − 1. Then *m*_2_ minimizes *h*(*u*) (p.0)

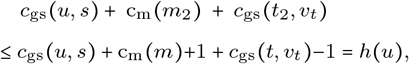

and furthermore *c*_seed_(*u, m*_2_) = *c*_seed_(*u, m*) (p.1), *c*_gap_(*u, m*_2_) = *c*_gap_(*u, m*) (p.2), and *c*_gap_(*m*_2_, *v*_*t*_) < *c*_gap_(*m, v*_*t*_) (p.3), contradicting the choice of *m*, so *c*_gap_(*t, v*_*t*_) > *c*_seed_(*t, v*_*t*_) is impossible and *t* ⪯_*T*_ *v*_*t*_.

#### Theorem 5

*Given a* consistent *set of matches* M, *the gap-chaining seed heuristic can be computed using scores in the transformed domain:*

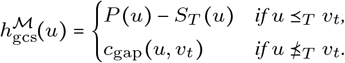

*Proof*. Write 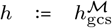 and 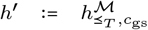. When 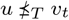, *h*(*u*) = *c*_gap_(*u, v*_*t*_) by Lemma 7. Otherwise, when *u*⪯_*T*_ *v*_*t*_, we have *h*(*u*) = *h*^′^(*u*). Let *u*⪯_*T*_ *m*_1_⪯_*T*_ … ⪯_*T*_ *v*_*t*_ be a ⪯_*T*_ -chain for *h*^′^ as in Def. 1. All terms in *h*^′^ satisfy end(*m*_*i*_) ⪯_*T*_ start(*m*_*i*+1_), so *c*_gap_ ≤ *c*_seed_ and by Lemma 6 *c*_gs_(*m*_*i*_, *m*_*i*+1_) = *c*_seed_(*m*_*i*_, *m*_*i*+1_). Thus, 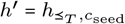 and 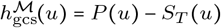 by Thm. 4.

## C Further results

### C.1 A*PA parameter grid search

We ran a parameter grid search over various *r* and *k* with 5 parallel jobs for both synthetic data and human data.

#### Synthetic data

Figs. 8a and 8b compare the runtime of A*PA with GCSH with DT for various *k* for both exact and inexact matches. For low divergence, exact matches are faster since they have a sufficiently large potential and do not require the overhead of finding all inexact matches, while for large divergence, the additional potential of inexact matches is required. In both cases, *k*=15 is a reasonable choice.

**Fig. 8.**
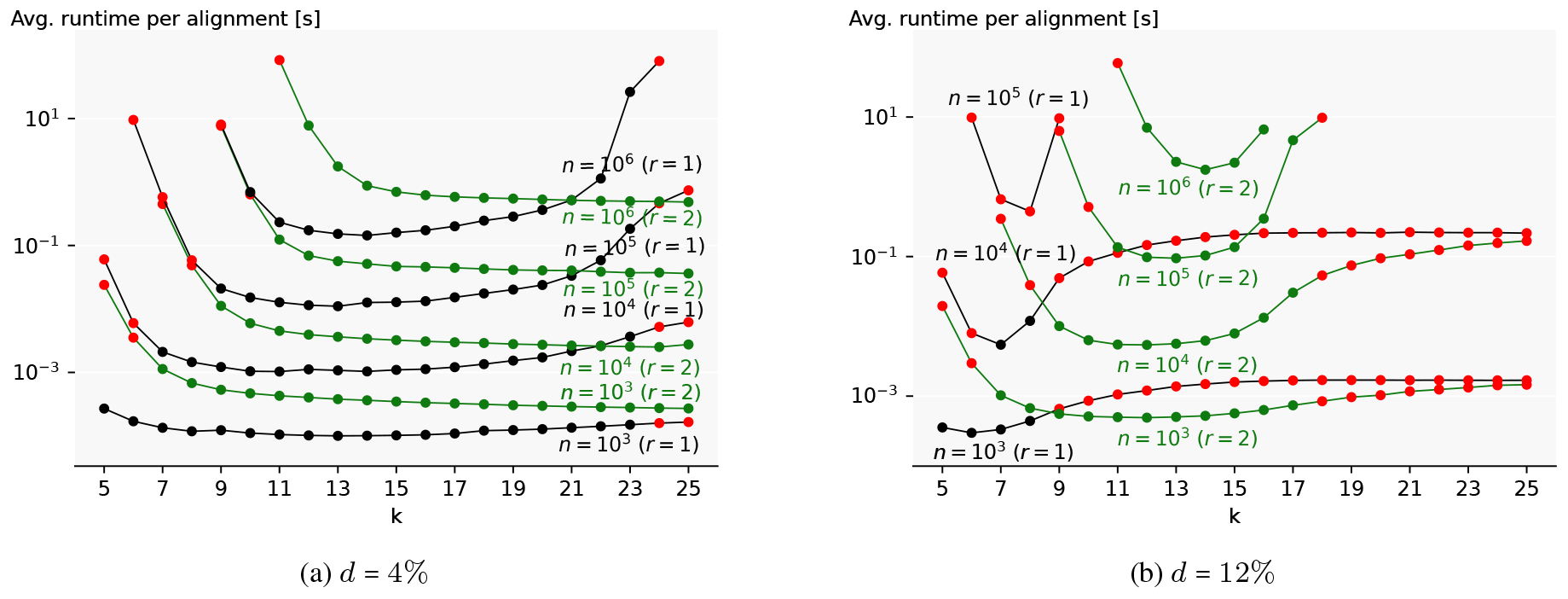
Parameter grid search on synthetic data. (*N* = 10^7^). Runtime of A*PA with GCSH with DT for varying *k* and *r* at *n* ∈ {10^3^, 10^4^, 10^5^, 10^6^ } bp and (a) *d*=4% and (b) *d*=12%. Black lines indicate *r*=1 and green lines indicate *r*=2. Missing datapoints are either due to timing out at *n*/*N* ⋅1000 s, or in 2 cases due to exceeding the 32 GiB memory limit. Red dots indicate alignments where *k* is too small or large (see App. C.1).

#### Human data

Fig. 9 shows that *r*=2 and *k*=15 minimize the runtime of the third quartile on both datasets. On the ONT reads without genetic variation, choosing a larger *k* slightly reduces the median runtime, but increases the number of timeouts, making *k*=15 a reasonable tradeoff.

**Fig. 9.**
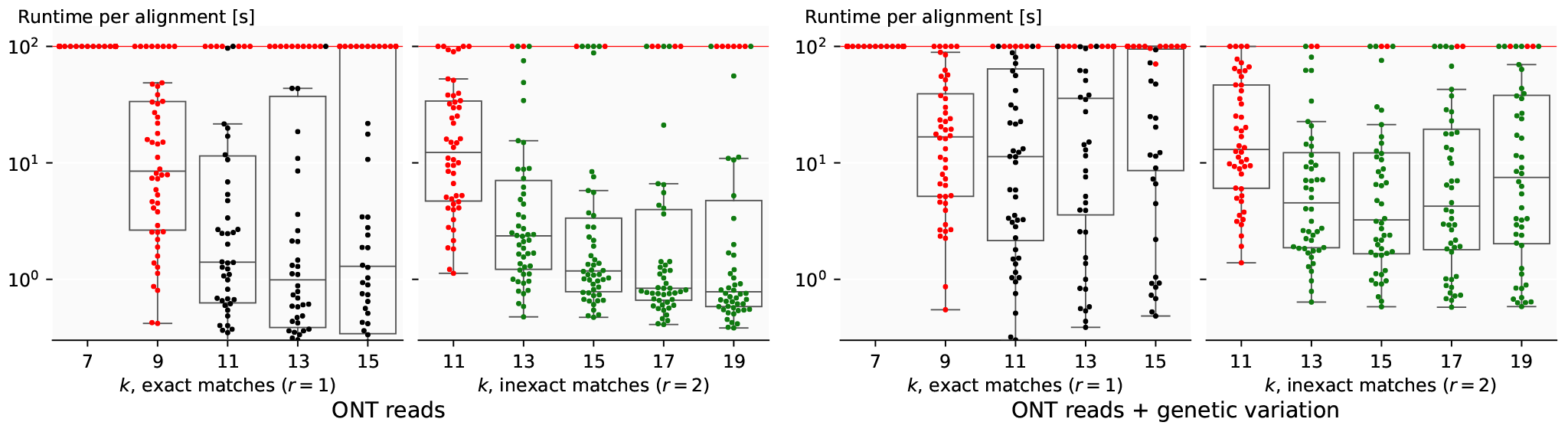
Parameter grid search on human data. (logarithmic). Plots show the runtimes on human ONT reads for varying *k* and *r* for A*PA with GCSH with DT. ONT reads are without (left) and with (right) genetic variation. Each dot shows the runtime for aligning a single sequence pair (capped at 100 s). Red dots indicate alignments where *k* is too small or large (see App. C.1).

#### Bounds on *k*

The runtime is generally not very sensitive to the exact choice of *k* as long as it falls within two bounds. First, too many spurious matches are prevented by setting *k* ≥ log_4_ *n* when *r*=1 and *k* ≥ 2 + log_4_ *n* when *r*=2. For example, *n*=10^6^ would require *k* ≥ 10 when *r*=1 or *k* ≥ 12 when *r*=2. Secondly, the potential *P* ≈ *r* ⋅*n*/*k* of the heuristic should be sufficiently larger than the divergence, which implies *k* should be a bit less than *r*/*d*. So, for *d*=12% we would choose *k* ≤ 1/0.12 = 8.3 when *r*=1 and *k* ≤ 2/0.12 = 16.7 when *r*=2.

Figs. 8 and 9 show alignments where *k* is not within these bounds in red, and indeed these alignments are relatively slow.

### C.2 Runtime scaling with length

In Fig. 4a we compare our A* heuristics with EDLIB and BIWFA in terms of runtime scaling with sequence length *n*.

### C.3 Expanded states and equivalent band

The main benefit of an A* heuristic is a lower number of expanded states, which translates to faster runtime. Instead of evaluating the runtime scaling with length (Fig. 10), we can judge how well a heuristic approximates the edit distance by directly measuring the *equivalent band* (Fig. 11) of each alignment: the number of expanded states divided by sequence length *n*, or equivalently, the number of expanded states per base pair. The theoretical lower bound is an equivalent band of 1, resulting from expanding only the states on the main diagonal.

**Fig. 10.**
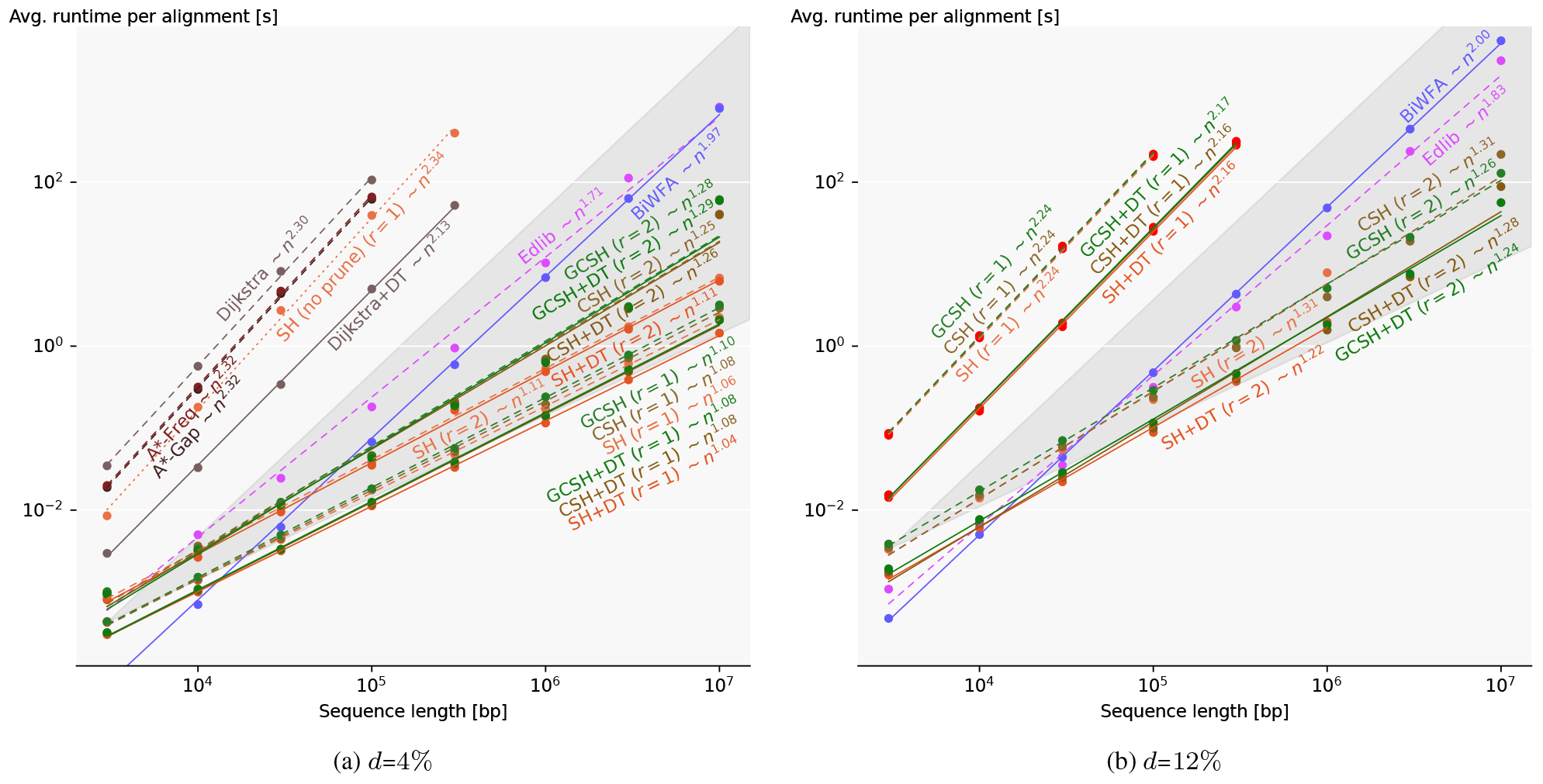
Runtime scaling with sequence length on synthetic data. Log-log plots comparing our heuristics with EDLIB and BIWFA. The slopes of the bottom (top) of the dark-grey cones correspond to linear (quadratic) growth. The seed length is *k*=15. Averages are over total *N* =10^6^ bp to *N* =10^7^ bp. SH without pruning is dotted, and variants with DT are solid. For *d*=12%, red dots show where the heuristic potential is less than the edit distance. Missing data points are due to exceeding the 32 GiB memory limit.

**Fig. 11.**
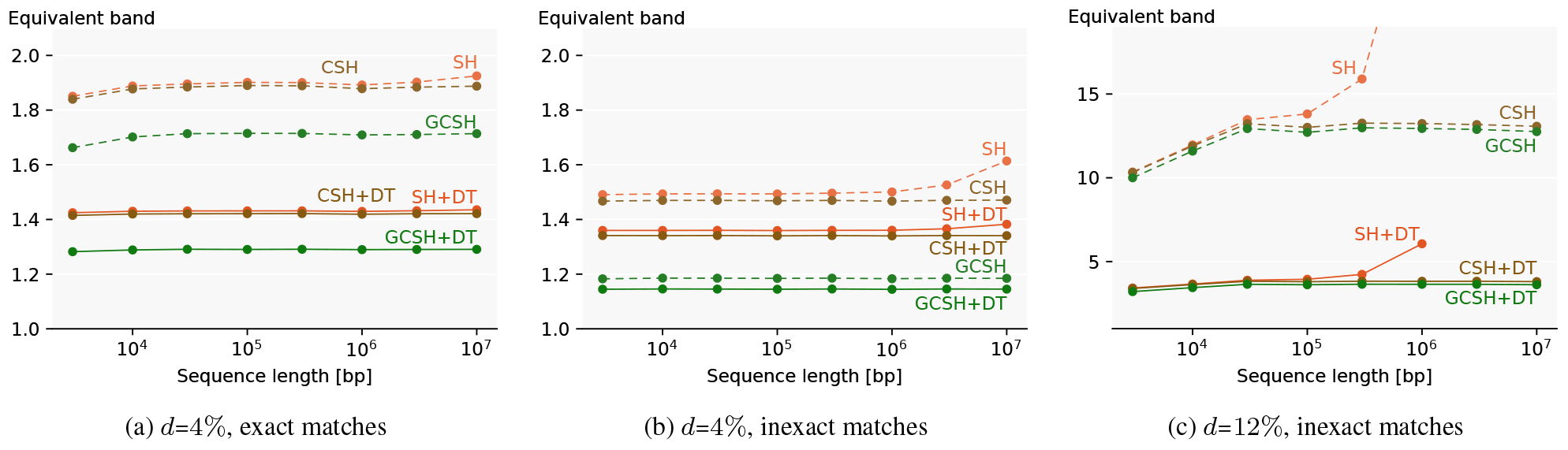
Equivalent band scaling with sequence length on synthetic data. (*k*=15). The equivalent band is the number of expanded states per bp for aligning synthetic sequences. Averages are over total *N* =10^7^ bp.

The equivalent band tends to be constant in *n*, indicating that the number of expanded states is linear on the given domain. The equivalent band of SH with inexact matches starts to grow around *n* ≥ 3 ⋅10^6^ at divergence *d*=4%, and around *n* ≥ 3 ⋅10^5^ at *d*=12%. Because of the chaining, CSH and GCSH cope with spurious matches and remain constant in equivalent band (i.e. linear expanded states with *n*). The equivalent band for GCSH is lower than CSH due to better accounting for indels. The DT variants expand fewer states by skipping non-farthest reaching states, also lowering the equivalent band.

### C.4 Runtime scaling with divergence: two modes

Figure 12 shows the runtime scaling with divergence for various heuristics. We notice two regimes of operation, depending on whether the heuristic potential *P* is sufficient to compensate for the edit distance: near-linear in *n* (constant in *d*) and quadratic in *n* (linear in *d*). The edit distance becomes larger than the potential *P* around *d* = *r*/*k*. For *k*=15 as in Fig. 12, the threshold is near *d*≈1/*k*=6.7% for exact matches and near *d*≈2/*k*=13.3% for inexact matches. Every error not accounted for by the heuristic triggers a search “to the side”, causing A* to explore *O*(*n*) additional states. When using DT, only *O*(*s*−*P*) additional farthest reaching states are explored instead, where *s* is the edit distance. This leads to observed runtimes of *O*(*n* + *n* ⋅max(*s* − *P*, 0)) without DT, and *O*(*n* + max(*s* − *P*, 0)^2^) with DT. These are similar to EDLIB’s *O*(*ns*) and BIWFA’s *O*(*n* + *s*^2^), but skipping over the first *P* errors.

**Fig. 12.**
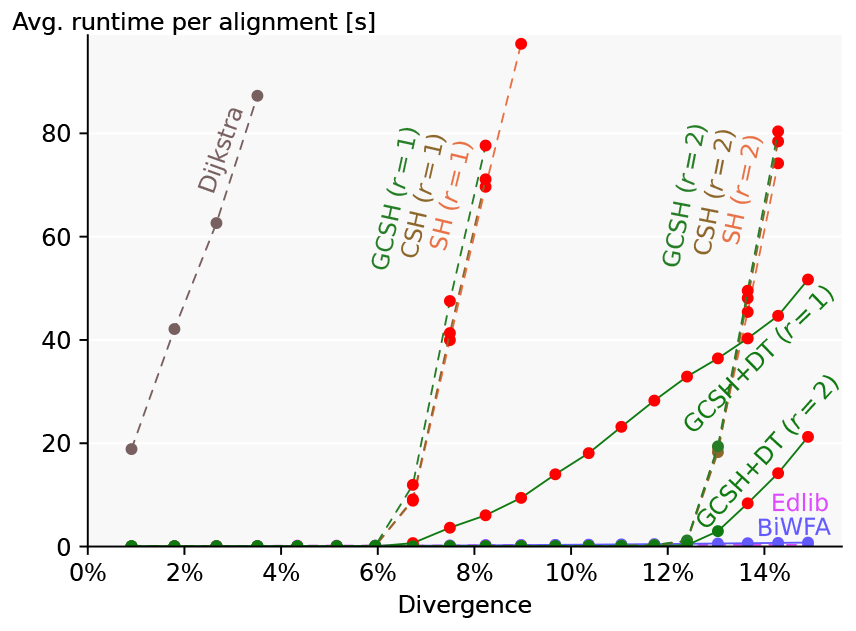
Runtime scaling with divergence. TODO LABELS (linear, synthetic, *n*=10^5^, 10^6^ bp total, *k*=15). The figure shows the same results as Fig. 4c, but zoomed out to show show scaling for high *d*, where runtime of A*PA degrades from constant in *d* to linear in *d. r*=1 and *r*=2 indicate exact and inexact matches, respectively. Red dots show where the heuristic potential is less than the edit distance. Missing datapoints timed out after 100 s.

### C.5 Human data statistic

Statistics on our human datasets are presented in Table 2.

**Table 2.** Human datasets statistics. ONT reads only include short gaps, while genetic variation also includes long gaps. **Cnt**: number of sequence pairs. **Max gap**: longest gap in the reconstructed alignment.

| Dataset | Cnt | Length [kbp] |  |  | Divergence [%] |  |  | Max gap [kbp] |  |  |
| --- | --- | --- | --- | --- | --- | --- | --- | --- | --- | --- |
|  |  | min | mean | max | min | mean | max | min | mean | max |
| ONT | 50 | 500 | 594 | 849 | 2.7 | 6.3 | 18.0 | 0.02 | 0.1 | 1 |
| ONT+gen.var. | 48 | 502 | 632 | 1053 | 4.4 | 7.4 | 19.8 | 0.05 | 1.9 | 42 |

### C.6 Memory usage

Table 3 compares memory usage of aligners on synthetic sequences of length *n*=10^6^. Alignments with inexact matches (*d*≥8%) use significantly more memory than those with exact matches because more *k*-mer hashes need to be stored to find inexact matches. Table 4 compares memory usage on the human data sets.

**Table 3.** Memory usage per algorithm. (synthetic data, *n*=10^6^). Exact matches are used when *d* ≤ 4%, and inexact matches when *d* ≥ 8%.

| Aligner | Memory usage [MB] |  |  |  |
| --- | --- | --- | --- | --- |
| | $d=1\%$ | $d=4\%$ | $d=8\%$ | $d=12\%$ |
| EDLIB | 2 | 1 | 2 | 2 |
| BiWFA | 15 | 13 | 14 | 18 |
| SH | 50 | 59 | 151 | 480 |
| CSH | 52 | 59 | 151 | 261 |
| GCSH | 49 | 50 | 151 | 255 |
| SH + DT | 49 | 49 | 151 | 150 |
| CSH + DT | 49 | 49 | 151 | 150 |
| GCSH + DT | 48 | 46 | 152 | 150 |

**Table 4.** Memory usage [MB] of aligners on human data. Medians are over all alignments; maximums are over alignments not timing out.

| Aligner | ONT reads |  | + genetic var. |  |
| --- | --- | --- | --- | --- |
|  | Median | Max | Median | Max |
| EDLIB | 2 | 5 | 2 | 6 |
| BiWFA | 11 | 19 | 15 | 24 |
| A*PA (GCSH + DT) | 160 | 3478 | 270 | 6926 |

### C.7 Runtime profile of A*PA

Figures 13a and 13b compare the time used by stages of A*PA. On synthetic data with exact matches (*r*=1), the runtime is spread over all parts of the algorithm. When using inexact matches, precomputation takes a significant fraction of the total time and updating contours becomes slower due to the increased number of matches.

**Fig. 13.**
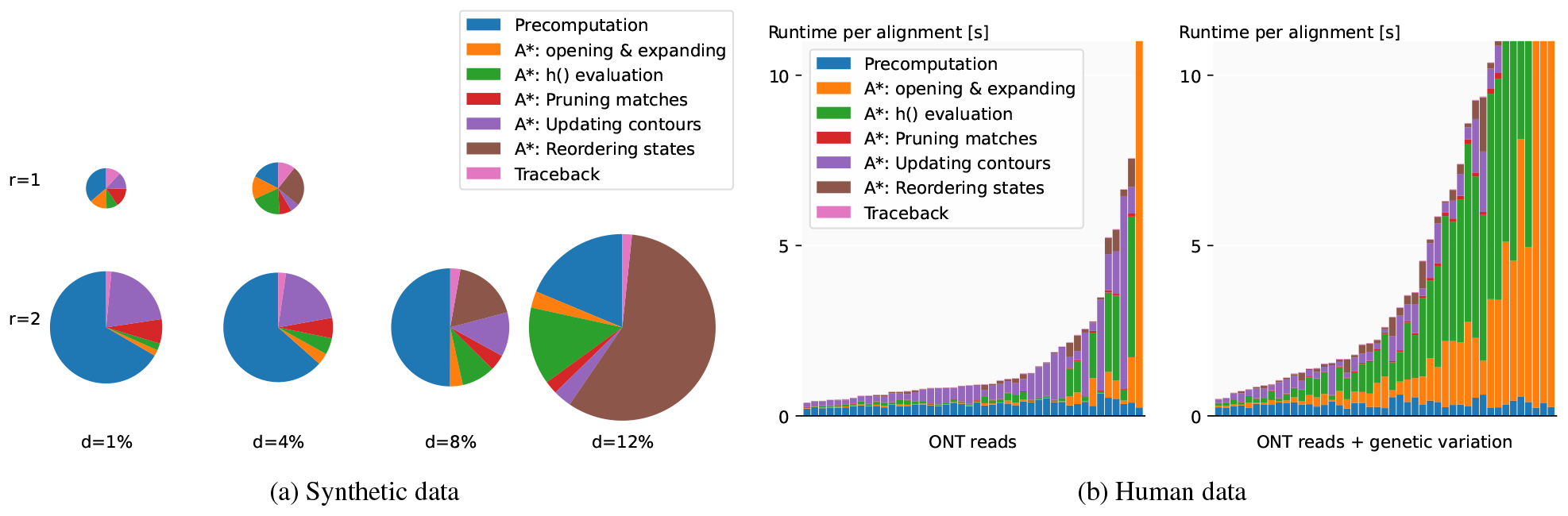
Runtime distributions per stage of A*PA (GCSH with DT) (stages do not overlap). Stage *A\** includes expanding and opening states. *Pruning matches* includes consistency checks. *Updating contours* includes updating of contours after pruning. (a) On synthetic data (*n*=10^6^ bp, *N* =10^7^ bp total, *k*=15). The circle area is proportional to the total runtime. Figures for *r*=1 and *d*≥8% are skipped due to timeouts (100 s). (b) On human data (*r*=2). Alignments are sorted by total runtime (timeouts not shown).

On human data, faster alignments spend a large fraction of their time on the precomputation, followed by the updating of contours after pruning matches. Slower alignments on the other hand are limited by the performance of the A* algorithm, and spend a large fraction of time on opening and expanding states, and evaluating the heuristic.

### C.8 Quadratic scaling in complex regions

Figure 14 shows the effect of complex regions in the sequences on GCSH (and thus on all our heuristics).

**Fig. 14.**
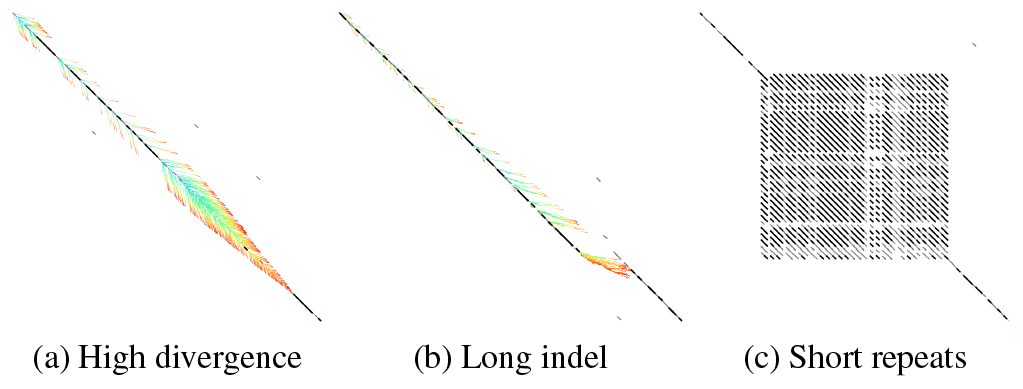
Quadratic exploration behavior for complex alignments. (GCSH with DT, *r*=2, *k*=10, synthetic sequences, *n*=1000). (a) A highly divergent region, (b) a deletion, and (c) a short repeated pattern inducing a quadratic number of matches. The colour corresponds to the order of expansion, from blue to red.

### C.9 Linear mode without matches

States are penalized by the number of remaining seeds that cannot be matched. So, curiously, matches are not always needed to direct the A* search to an optimal path. In fact, when each seed contains exactly one mutation seed heuristic scales linearly even though there are no matches, as shown by the artificial example in Fig. 15.

**Fig. 15.**
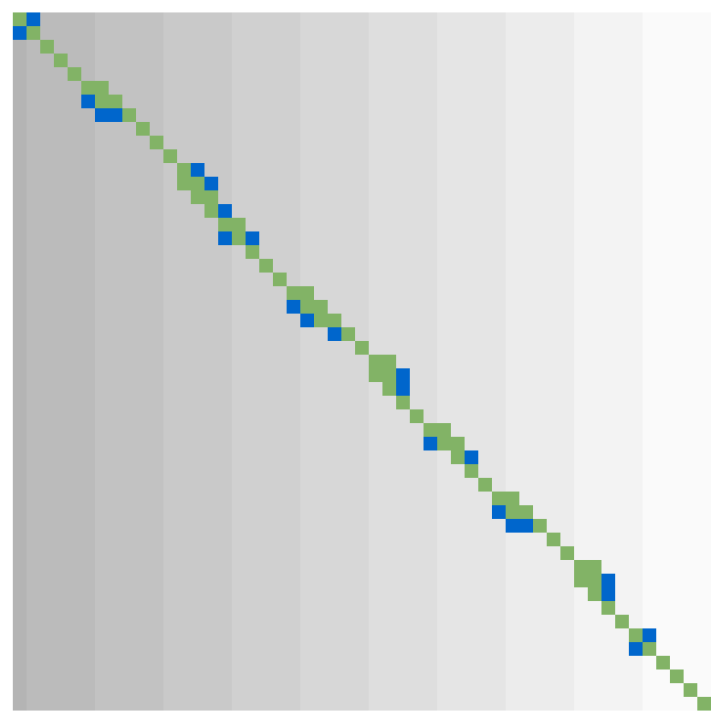
Artificial example of A* with seed heuristic with no matches. (*n*=*m*=50, *r*=1, *k*=5, 80% similarity, 1 mutation per seed alternating substitution, insertion, and deletion). The background colour indicates *h*_s_(*u*) with higher values darker. Expanded states are green 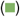, open states blue 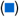.

### C.10 Comparison of heuristics and techniques

Figure 16 shows the effect of our heuristics and optimizations for aligning complex short sequences. The effect of pruning is most noticeable for CSH and GCSH without DT. GCSH is our most accurate heuristic, so, as expected, it leads to the lowest number of expanded states.

**Fig. 16.**
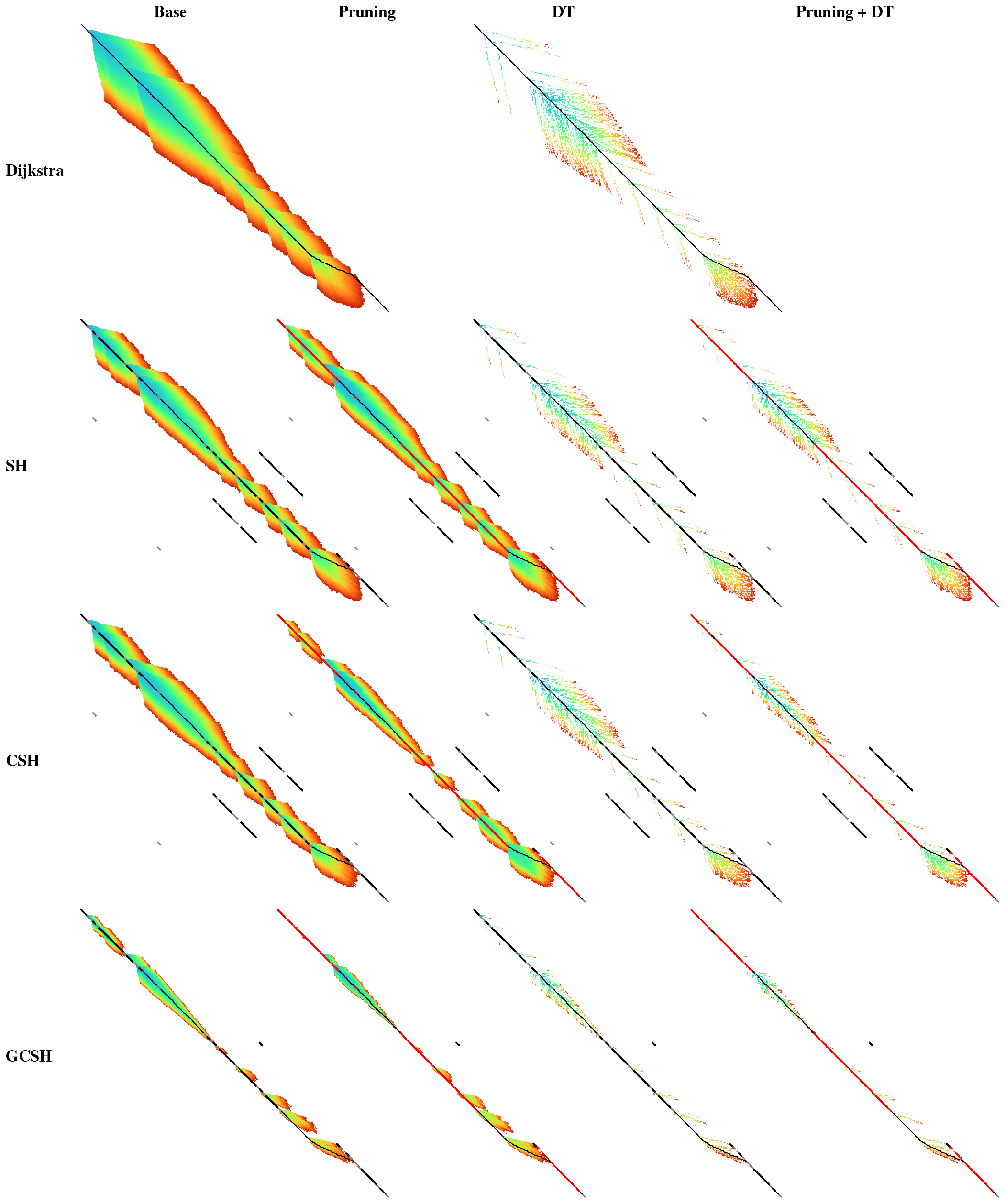
Expanded states for various heuristics and techniques, on a sequence containing a noisy region, a repeat, and an indel. (*n*=1000, *d*=17.5%). The colour shows the order of expanding, from blue to red. The sequences include a highly divergent region, a repeat, and a gap. Matches are shown as **black diagonals**, with inexact matches in **grey** and pruned matches in red. The final path is **black**. Dijkstra does not have pruning variants, and Dijkstra with DT is equivalent to WFA. More accurate heuristics reduce the number of expanded states by more effectively punishing repeats (CSH) and gaps (GCSH). Pruning reduces the number of expanded states before the pruned matches, and diagonal transition reduces the density of expanded states in quadratic regions.

## D Notation

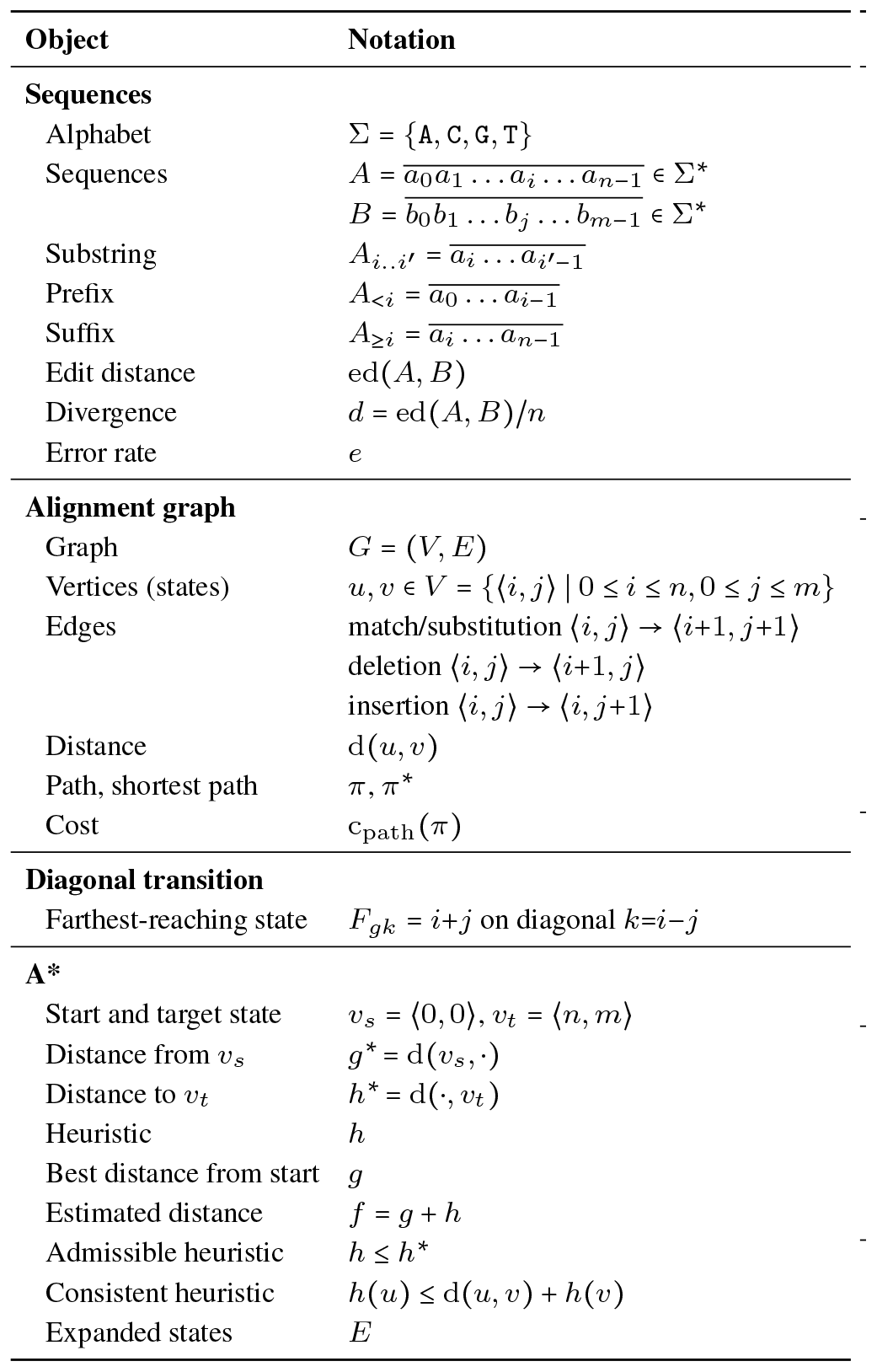

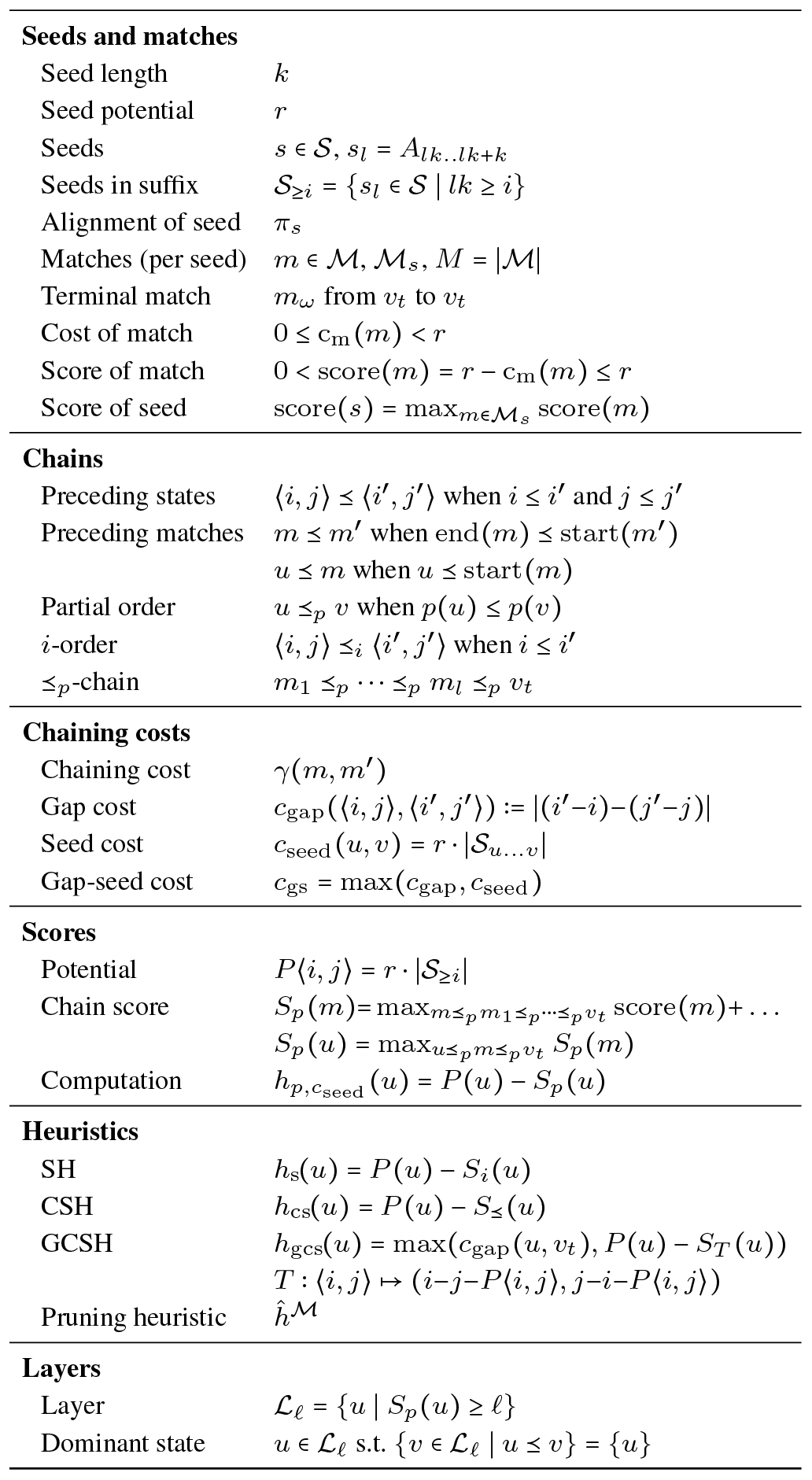

## Footnotes

1 github.com/RagnarGrootKoerkamp/astar-pairwise-aligner (tag [evals](https://github.com/RagnarGrootKoerkamp/astar-pairwise-aligner/releases/tag/evals))

2 github.com/pairwise-alignment/pa-bench/releases/tag/datasets

3 github.com/pairwise-alignment/pa-bench (tag astarpa-evals)

4 Previous works indicate the column *i* of *u*, but using the antidiagonal *i*+*j* keeps the symmetry between insertions and deletions.

